# Systematic identification of pH-sensing amyloid core motifs reveals a widespread mechanism for reversible protein assembly upon stress

**DOI:** 10.64898/2026.08.11.744147

**Authors:** Anastasiia Kovalenko, Dorota M. Pfizenmaier, Caroline Wilson-Zbinden, Federico Uliana, Izabella Krystkowiak, Martina Bonassera, Claudia C. Schmidt, Pavel Afanasyev, Tamino Cairoli, Alvar D. Gossert, Tanushree Agarwal, Sonja Kroschwald, Viviane Lutz Bueno, Gea Cereghetti, Tuomas Knowles, Norman Davey, Matthias Peter

## Abstract

Unlike irreversible pathological amyloids, reversible fibrils can be regulated via pH-sensing core motifs, characterized by amyloid properties unleashed upon stress-induced protonation of critical residues. Here, we combined bioinformatic predictions and an *in vitro* validation pipeline to search for pH-responsive, reversible amyloid core peptides in yeast and human proteomes. This approach uncovered biophysical properties distinguishing pH-sensing and constitutive amyloid cores and established reliable criteria to identify novel reversible assemblies based on sequence data. Selected full-length candidate proteins with evolutionarily conserved pH-sensing motifs were analyzed in *Saccharomyces cerevisiae* using fluorescence microscopy and SDS-resistance assays, revealing multiple proteins forming reversible assemblies in stationary phase. We found that protonation of a specific histidine in the amyloid core motif of the asparagine synthase Asn1 is necessary and sufficient for assembling catalytically inactive, reversible structures called cytoophidia. Interestingly, mutant cells that fail to assemble Asn1-cytoophidia show defects to recover from stationary phase, demonstrating functional relevance of pH-sensing amyloid core motifs *in vivo*. Taken together, we uncovered a widespread and conserved pH-sensing mechanism that regulates the reversible assembly and function of structurally diverse fibrils upon stress.

## Introduction

Protein or peptide aggregation into amyloid fibrils is characterized by the formation of cross b-sheet structures (Nelson *et al*., 2005; Dobson, Knowles and Vendruscolo, 2020; Ke *et al*., 2020). This stable amyloid fold is thought to disrupt cellular functions and ultimately affect viability, causing neurodegenerative disorders like Parkinson’s and Alzheimer’s disease (Knowles, Vendruscolo and Dobson, 2014). Yet, recent work discovered the existence of functional amyloids, which can switch between their native and fibrillar states in a reversible and regulated manner (Fowler *et al*., 2007; Otzen and Riek, 2019). Reversible amyloids are part of a cellular stress response to nutrient starvation or heat shock, and they play critical roles in processes such as metabolism, hormone release and long-term memory (Maji *et al*., 2009; Majumdar *et al*., 2012; Saad *et al*., 2017). Fundamental questions concern the structural distinctions between pathological and reversible amyloid fibrils, the underlying assembly and disassembly regulatory mechanisms, and how functional amyloid fibrils avoid cellular toxicity (Jackson and Hewitt, 2017). It is also unclear whether chronic stress conditions or genetic mutations may convert physiological fibrils into toxic amyloids.

Functional fibrils may be controlled by post-translational modifications, metabolite binding and dedicated chaperone systems. For example, phosphorylation of the yeast RNA-binding protein Rim4 by Ime2 modulates its disassembly, ensuring progression through meiosis (Carpenter *et al*., 2018). Moreover, binding of the allosteric activator fructose-1,6-bisphosphate promotes disassembly of yeast pyruvate kinase (PK) Cdc19 amyloids, which is further enhanced by the disaggregase Hsp104 (Cereghetti *et al*., 2021). Finally, oxidation of specific cysteine residues triggers amyloid formation of human IL-38 (Diaz-Barreiro *et al*., 2024) and the W protein of Nipah and Hendra viruses (Gondelaud *et al*., 2024).

Importantly, reversible aggregation of several proteins is regulated by intracellular pH changes in a variety of physiological and pathological conditions. Examples include human peptide hormones, the translation regulator CPEB4, and PK Cdc19 and its human homolog PKM2 (Maji *et al*., 2009; Cereghetti *et al*., 2024; Garcia-Cabau *et al*., 2025). Amyloid conversion of PK upon starvation or heat stress inhibits catalytic activity and may prevent stress-induced degradation. Once the stress is overcome, PK fibrils rapidly disassemble, enabling rapid restart of ATP production by converting phosphoenolpyruvate (PEP) into pyruvate (Saad *et al*., 2017; Cereghetti *et al*., 2021). Indeed, cytosolic pH drops upon heat stress, ageing and starvation in yeast (Imai and Ohno, 1995; Joyner *et al*., 2016; Knieß and Mayer, 2016; Munder *et al*., 2016; Triandafillou *et al*., 2020), while in humans low cytosolic pH correlates with neurodegenerative diseases (Fang et al., 2010; Schwartz et al., 2020). In contrast, cancer cells retain elevated intracellular pH (Webb *et al*., 2011), which enhances glycolysis and promotes proliferation (Koch et al., 2020). However, despite the widespread importance, the global effects of intracellular pH changes on protein aggregation remain poorly understood.

pH-sensing has been studied in Cdc19 and was shown to be mediated by a short motif referred to as amyloid core, which becomes protonated in response to stress conditions, causing its transition into an amyloid state (Cereghetti *et al*., 2024). Likewise, protonation of a histidine (H391) in the PKM2 amyloid core triggers reversible fibril formation, while the corresponding residue in another human PK isoform - PKM1 - is flanked by charged amino acids that prevent amyloid conversion. Therefore, pH-sensing by dedicated amyloid core motifs may serve as a conserved mechanism for inducing reversible amyloid formation, particularly upon cellular stress. Yet, the efforts to study pH-sensing core motifs are at early stages, and their functional relevance and biochemical properties remain elusive.

Structural methods such as circular dichroism (CD), Fourier transform infrared spectroscopy (FTIR), electron microscopy (EM), atomic force microscopy (AFM), or light scattering techniques report specific protein folds and conformations. Despite recent advances (Scheres, Ryskeldi-Falcon and Goedert, 2023; Garcia-Pardo and Ventura, 2024), throughput and sample requirements are generally limiting for discovery of new functional amyloids or other types of fibrils (Nilsson, 2004; Li *et al*., 2009). Measuring amyloid properties thus often involves conformation-sensing dyes such as classically used Thioflavin T (ThT) or Congo Red (CR) (Dirvy, 1927; Vassar and Culling, 1959), or testing structural robustness with detergent resistance, for example with semi-denaturing detergent agarose gel electrophoresis (SDD-AGE) (Halfmann and Lindquist, 2008). In practice, a combination of different methods is typically used to characterize amyloid structures (Matiiv *et al*., 2020).

A bottleneck to identify functional amyloids has been the lack of robust predictive and experimental tools. Powerful amyloid prediction algorithms have been developed, but they were mostly trained on classical pathological or artificial amyloids (Fernandez-Escamilla *et al*., 2004; Zhang, Chen and Lai, 2007; Zibaee *et al*., 2007; Tian *et al*., 2009; Maurer-Stroh *et al*., 2010). Recent identification of functional fibrillar assemblies with amyloid properties that do not have cross-β-sheet folds (Tayeb-Fligelman *et al*., 2017; Dey *et al*., 2024) has prompted reconsidering the amyloid definition (Benson *et al*., 2018; Matiiv *et al*., 2020; Buxbaum *et al*., 2022). Indeed, functional, reversible amyloids might display structural and biochemical heterogeneity that is not adequately captured by current bioinformatic tools.

To circumvent these restrictions, we developed an *in silico* workflow to predict pH-sensing amyloid core motifs, coupled to a robust *in vitro* validation method based on short peptides. By applying these tools to the human and *S. cerevisiae* proteomes, we identify novel amyloid core motifs and deduce hallmarks distinguishing pH-sensing, reversible and constitutive assemblies. We combine biochemical and cell-based analysis to study selected yeast candidates conserved in human cells, among them asparagine synthetase Asn1. Indeed, upon starvation, Asn1 forms elongated assemblies termed cytoophidia (Zhang *et al*., 2018), and we show that these reversible structures require protonation of a histidine located in a conserved amyloid-core motif. Together, this work reveals that conserved pH-sensing amyloid core motifs can assemble different types of reversible fibrils and are part of a widespread mechanism to regulate enzyme activity during stress.

## Results

### Bioinformatic criteria and iterative in vitro validation define pH-sensing amyloid core motifs

To systematically identify reversible pH-sensing amyloids, we set up a bioinformatic screen coupled to an experimental validation pipeline which we called ‘<u>T</u>hioflavin T <u>A</u>nd <u>P</u>elleting <u>A</u>ssays for <u>A</u>myloid <u>S</u>creening’ (TAPAAS) with 15-mer peptides (Fig. 1A). Initial bioinformatic criteria were defined by the non-amyloid PKM1 and pH-sensing amyloid PKM2 core motifs (Cereghetti *et al*., 2024) (Fig. S1A). Both motifs contain aromatic and hydrophobic residues, and a protonatable histidine. PKM1 has 2 positively charged residues following the histidine (H-R-K), whereas the PKM2 histidine is followed by neutral residues (H-L-Q). First, SLiMSearch (Krystkowiak and Davey, 2017) was used to identify 15-mer peptides with a central histidine within intracellular human proteins. Proline-containing peptides were excluded due to their known disruptive effect on β-sheet formation (Gurung *et al*., 2023). Then we annotated peptides as pH-sensing with the following criteria. (1) Peptides contained at least two hydrophobic and/or aromatic residues, with at least one located adjacent to the central histidine; (2) No more than 2 charged amino acids with the same charge were allowed to cluster in the sequence, as clustered charges create repulsive forces that disfavor amyloid formation. (3) Peptides were predicted to have amyloid properties when we applied the AMYLPRED2 tool using both the default threshold (score) and an arbitrary consensus coverage criterium (Tsolis *et al*., 2013). For the TAPAAS pipeline we used two experimental strategies to validate reversible amyloid formation. In the first, 15-mer synthetic peptides were measured for Thioflavin T (ThT) binding incubated at either pH 5.0 or 7.5, and conversely, amyloid reversibility was assessed by pre-incubating the peptides at pH 5.0 and then shifting them to pH 7.5 or maintaining pH 5.0 for control. In the second, we established a pelleting assay combined with mass spectrometry readout that quantitatively reports the extent of aggregation (Fig. 1A). To control the TAPAAS workflow we used peptides encompassing the 15-mer core motifs of PKM1 and PKM2 (Fig. S1B).

**Figure 1:**
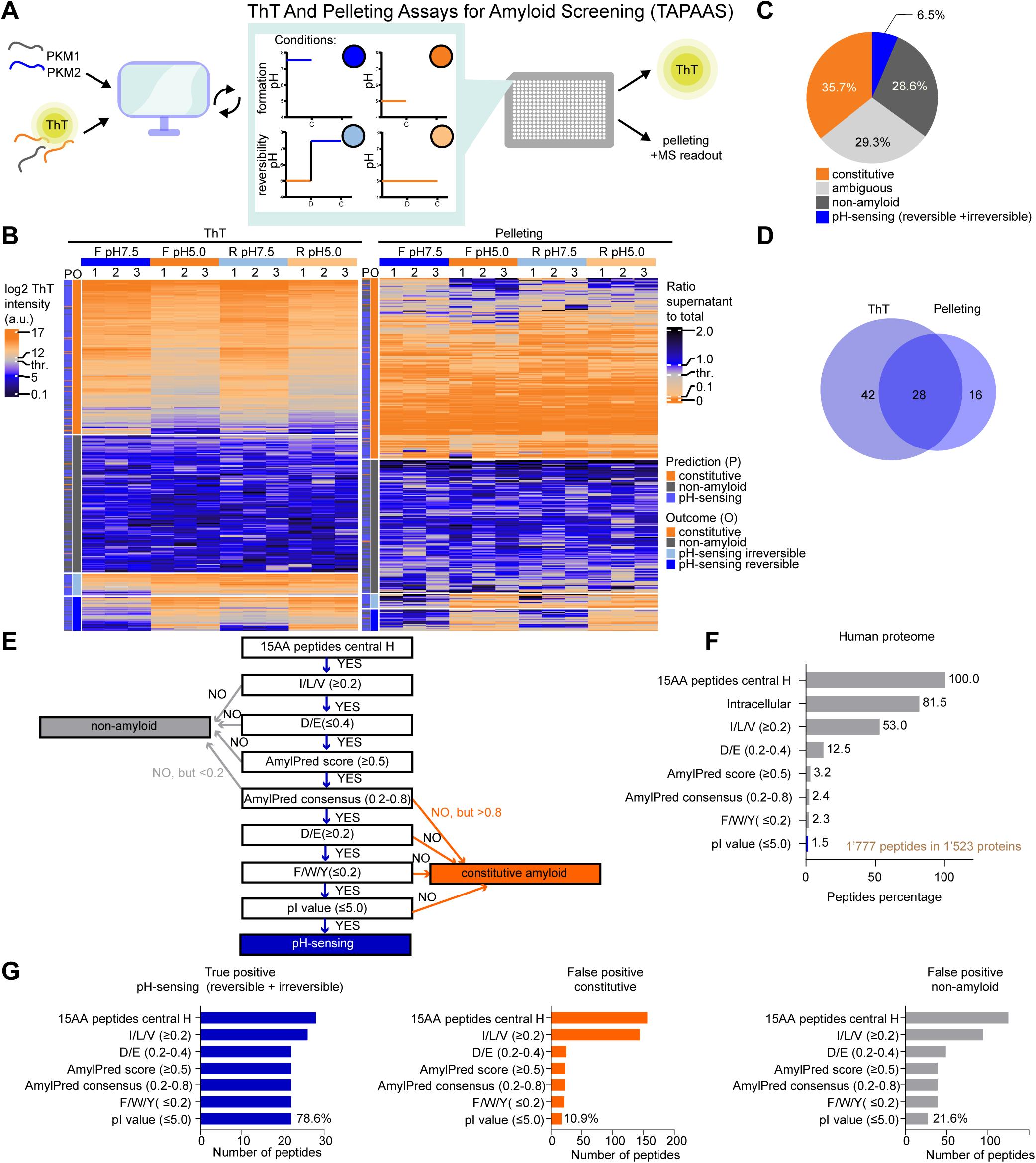
Bioinformatic criteria and iterative *in vitro* validation define pH-sensing amyloid core motifs. A. Schematic representation of the screening method combining bioinformatic predictions with a peptide-based *in vitro* validation pipeline ThT And Pelleting Assays for Amyloid Screening (TAPAAS). The experimental workflow probes the formation and reversibility of amyloids at different pH conditions. Iterative cycles were used to improve the bioinformatic screening parameters, initially set by the core motif properties of non-amyloid PKM1 and pH-sensing, reversible amyloid PKM2. D - sample dilution to change pH, C - sample collection for measurements. B. Heat maps representing TAPAAS sub-assays, ThT binding (left) and pelleting (right), results for 437 synthetic 15-mer peptides. Amyloid formation (F) and reversibility (R) were tested at pH 7.5 or 5.0. Three experimental replicates are plotted in separate columns (1 – 3). Predictions (P) and outcome (O) classification are indicated in the columns on the left; a.u.: arbitrary units; thr: arbitrary amyloid thresholds were set to 500 a.u. for ThT intensity, and 0.715 ratio of supernatant to total for pelleting assay. C. Pie-chart summarizing the TAPAAS outcome for 437 analyzed synthetic 15-mer peptides with a central histidine. The screen identified 28 (6.5 %) peptides assembling pH-sensing amyloids, 125 (28.6%) non-amyloids and 156 (35.7%) constitutive amyloids. Peptides classified as “ambiguous” (n=128, 29.3%) include those with results in two TAPAAS sub-assays that were not coherent. D. Venn diagram showing the number of pH-sensing peptides with overlapping or distinct outcome of the two TAPAAS sub-assays. E. Schematic decision map for categorizing predicted amyloid core peptides, using the criteria and parameters improved by the TAPAAS outcome. The workflow distinguishes pH-sensing, constitutive- and non-amyloid-forming peptides. F. The bioinformatic workflow was applied to the complete human proteome, and total 15-mer peptides with a central histidine and no prolines is represented as 100%. The percentage (%) of retained peptides after sequentially applying the selection criteria is plotted, resulting in a total of 1’777 peptides (1.5%) among 1’523 proteins that qualify as pH-sensing amyloid core candidates. G. Bar plot showing the performance of the filtering criteria from panels E and F on pH-sensing amyloids (n=28), constitutive amyloids (n=156) and non-amyloid peptides (n=125) identified by the TAPAAS assay. The resulting true positive (TPR) and false positive (FPR) rates are indicated.

Experimental validation among 51 initial candidates detected 7 pH-sensing peptides that displayed significantly higher ThT intensity at low pH compared to near-neutral pH of 7.5 (Fig. S1C, Table S1). Some candidates were also reversible, displaying significantly lower ThT intensity after shifting the pH back to the near-neutral pH versus the low pH-control (Fig. S1D). Since these pH-sensing amyloid core peptides were characterized by a low isoelectric point (pI) value (Fig. S1E), we refined the bioinformatic search criteria by adding a pI value constraint. This search predicted 6’519 pH-sensing amyloid cores from 4’234 different human proteins (Fig. S1F). Of these, we chose 306 pH-sensing test peptides. Moreover, we selected 198 control peptides that violated either single or multiple filtering criteria and were thus predicted to be either non-pH-sensing constitutive amyloids (n=22) or non-amyloids (n=176). Among these 504 peptides (Table S2) analyzed via TAPAAS, 67 were excluded due to issues with peptide synthesis or mass spectrometry (MS) identification and quantification. For the remaining peptides, 70% showed coherent outcome between pelleting and ThT binding assay resulting in 309 unambiguously classified peptides: 125 non-amyloid, 156 constitutive amyloid and 28 reversible or irreversible pH-sensing amyloid cores (Fig. 1B and C, D, Table S2, S3). Based on this TAPAAS outcome the bioinformatic predictions with the pI constraint (Fig. S1F) were only 34% accurate (Table S3).

To improve the bioinformatic criteria, we compared the biophysical properties of these newly identified reversible amyloid peptides to constitutive and non-amyloid forming counterparts and examined the amino acid frequencies in the pH-sensing amyloid cores (Fig. 2). Incorporating the experimental data, we then further adjusted the filtering criteria (Fig. 1E) to (1) specify the required content of hydrophobic, aromatic and negatively charged amino acids within the core, (2) lower the pI value threshold, (3) remove the ineffective same-charge clusters exclusion, and (4) relax the AMYLPRED2 consensus coverage criterion. This optimized algorithm predicted 1’777 peptides from 1’523 human proteins, a 3-fold reduction of potential hits compared to the original conditions (Fig. 1F, Table S4). When applying this advanced algorithm to the validated TAPAAS results, we observed a ∼2-fold improvement in accuracy to 66%. Moreover, the false positive rate (FPR) was reduced 8-fold to 11% for constitutive and 2-fold to 22% for non-amyloid peptides, and the true positive rate (TPR) was almost 80% (Fig. 1G, Table S5). To substantiate these newly predicted peptides from the human proteome (Fig. 1F), we validated a random subset by ThT binding, and indeed, 68% were classified as pH-sensing with the large majority at least partially reversible (Fig. S1G-H, Table S1). Hence, the bioinformatic pipeline with experimentally validated parameters improved the precision almost 7-fold compared to the initial set-up, making the workflow a powerful tool to identify potential pH-sensing, reversible amyloids in any organism with a sequenced genome.

**Figure 2:**
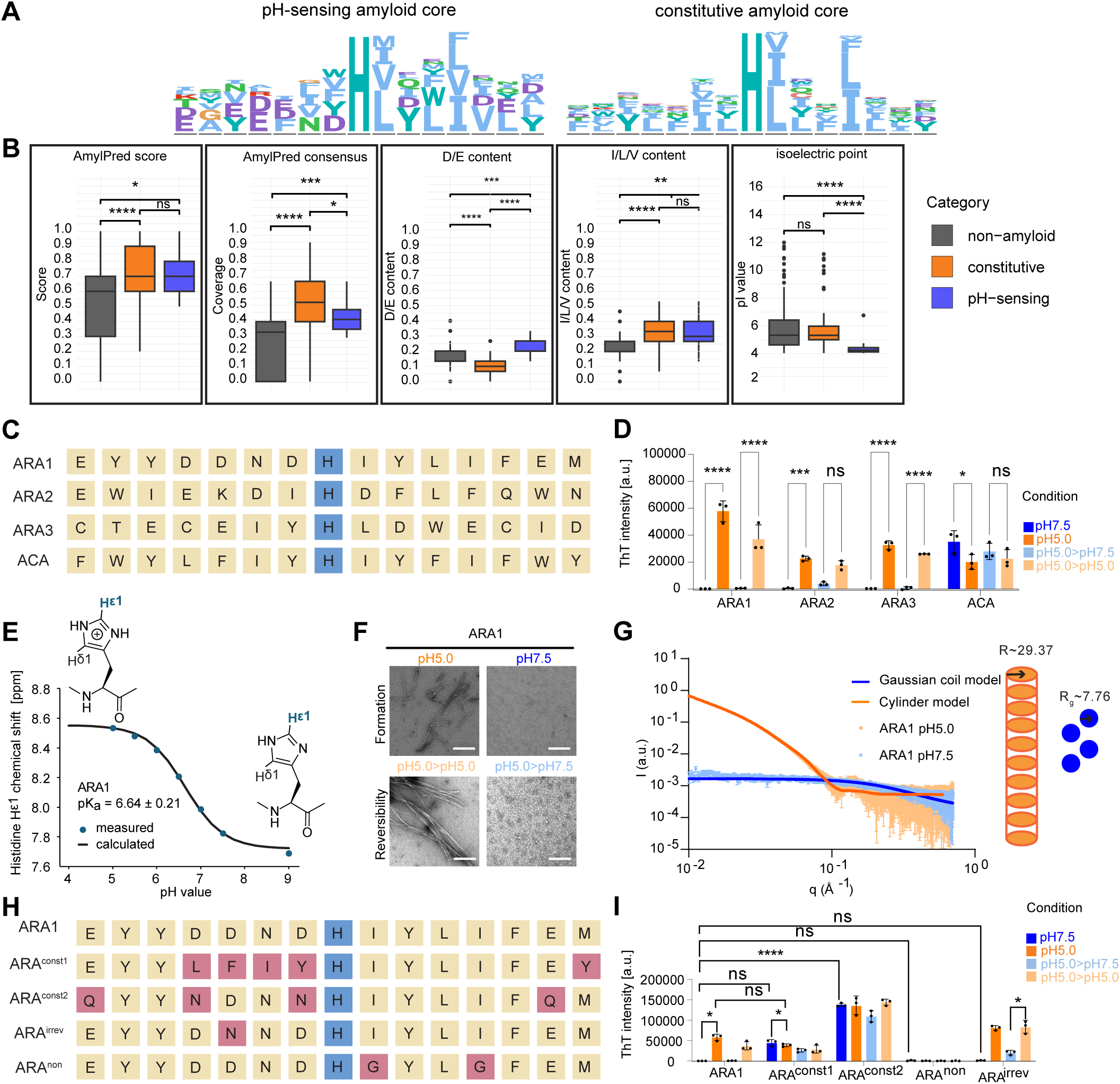
Biophysical properties of identified amyloid cores allow for rational design of artificial peptides. A. Sequence logos of TAPAAS validated 15-mer peptides with a fixed central histidine (H) forming pH-sensing (left, n=28) or constitutive amyloids (right, n=156). B. Comparison of selected properties including acidic (D/E) and specific hydrophobic (I/L/V) amino acid content, isoelectric point (pI) value, AMYLPRED2 score and consensus coverage of experimentally validated pH-sensing (n=28), constitutive (n=156) and non-amyloid (n=125) peptides. The box plots show median, Q1-Q3 ± max 1.5 IQR and outliers, Kruskal-Wallis and Dunn post-hoc-test with Holm correction; * p <0.05, ** p <0.01, *** p <0.001, **** p <0.0001; ns, not significant. C. Sequence representation of three predicted <u>a</u>rtificial pH-sensing, reversible <u>a</u>myloids (ARA1 - 3). For control, a predicted <u>a</u>rtificial <u>c</u>onstitutive <u>a</u>myloid (ACA) was included. D. ThT binding of ARA1, ARA2, ARA3, and ACA peptides analyzed at the indicated buffer conditions. For formation, peptides were incubated for 24 h at 30° at pH 7.5 or pH 5.0. To probe reversibility, the peptides were pre-incubated at pH 5.0 for 24 h at 30°C and then maintained at pH 5.0 for control or shifted to pH 7.5 for an additional 24 h at 30°C; n=3, mean ± SD; a.u.: arbitrary units. A two-way ANOVA test, followed by Tukey’s multiple-comparisons controls statistical significance; * p <0.05, *** p <0.001, **** p <0.0001; ns, not significant. E. Determination of the pKₐ value of the central histidine in the ARA1 peptide by nuclear magnetic resonance (NMR) spectroscopy. Chemical shift changes (data points) were monitored across a pH gradient from 4 to 9, yielding a fitted pKₐ of 6.64 ± 0.21. F. Representative negative-staining transmission electron microscopy (ns-TEM) of the ARA1 peptide treated as in panel D to allow formation of fibrils (top row) and test dissolution (bottom row); n=3, scale bar: 200 nm. G. ARA1 peptide was analyzed by small-angle X-ray scattering (SAXS) after 24 h at 30°C to allow formation of fibrils at pH 5.0 or pH 7.5. SAXS data were visualized as log-log plot of the scattering intensity (I) versus the scattering vector (q). At pH 7.5, the ARA1 peptide remains as unfolded monomers (schematic), fitted with a Gaussian coil model. In contrast, at pH 5.0, the ARA1 peptide exhibits scattering as large, elongated structures, which is well-modelled as a cylinder with an elliptical cross-section (schematic). Data are representative of n=4 (dotted lines) and fitted to the corresponding models (solid lines). H. Sequence representation of designed ARA1 mutant peptides, with the changed amino acids highlighted in red. ARA^const1^ and ARA^const2^ are expected to form constitutive amyloids, ARA^irrev^ pH-sensing irreversible amyloids, and the changes in ARA^non^ are predicted to abolish amyloid formation at either pH. I. ThT binding of ARA1, ARA^const1^, ARA^const2^, ARA^non^ and ARA^irrev^. Peptides were treated as in panel D; n=3, mean ± SD; a.u.: arbitrary units. A two-way ANOVA test, followed by Tukey’s multiple-comparisons, controls statistical significance; * p <0.05, **** p <0.0001; ns, not significant; only relevant comparisons are shown.

### Biophysical properties of identified amyloid cores allow for rational design of artificial peptides

Comparing the biophysical properties of the validated peptide data allowed generating a binomial sequence logo of pH-sensing and constitutive amyloid peptides, highlighting the key properties (Fig. 2A). The most striking feature distinguishing pH-sensing amyloids from constitutive was the presence of multiple negatively charged residues, glutamic and aspartic acids (D/E content). These charged residues were enriched at the peripheral regions and contributed to their significantly lower pI-values compared to constitutive amyloids (Fig. 2B, S2A). It is possible that the expected repulsion forces are counteracted by protonation at low pH, in addition to protonation of the central histidine. In contrast, positively charged residues (K/R content) were underrepresented both in constitutive and pH-sensing amyloid cores, and constitutive amyloids showed significantly higher content of aromatic residues (F/W/Y content) compared to non-amyloid forming peptides. Both pH-sensing and constitutive amyloid cores had high content of hydrophobic residues (I/L/V content) and a high AMYLPRED2 score. The AMYLPRED2 consensus coverage was low, intermediate and high for non-amyloid, pH-sensing and constitutive amyloid cores, respectively. The IUPred analysis (Dosztányi *et al*., 2005) revealed that constitutive amyloid cores were significantly less disordered compared to non-amyloid controls, with pH-sensing peptides scoring in-between (Fig. S2A).

To experimentally probe the identified general properties (Fig. 2A), we engineered three <u>a</u>rtificial reversible <u>a</u>myloid peptides (ARA1, ARA2 and ARA3) and one <u>a</u>rtificial <u>c</u>onstitutive <u>a</u>myloid peptide (ACA), respectively (Fig. 2C, Table S1). Indeed, ThT binding assays demonstrated that the synthetic ARA peptides bound ThT at pH 5.0 but not at 7.5 displaying the expected pH-sensing properties, and ARA1 and ARA3 were reversible (Fig. 2D). In contrast, the synthetic ACA peptide bound ThT in all pH conditions (Fig. 2D). NMR measurements confirmed that the central histidine residue in the ARA1 peptide was protonated with a pKa value of 6.64 ± 0.21 (Fig. 2E, S2B), consistent with the observed amyloid conversion. A reduced overall intensity of the NMR signal was observed at lower pH, reflecting the decreased soluble peptide concentration due to aggregation (Fig. S2C). Circular dichroism (CD) measurements corroborated the ThT binding assays and indicated no defined structural properties of the ARA1 peptide at pH 7.5, but a strong negative peak at around 215 nm (Fig. S2D), characteristic of β-sheets (Rogers *et al*., 2019). This 215 nm peak disappeared again upon pH change to 7.5 (Fig. S2D), indicating reversibility of the assembly. Moreover, negative staining transmission electron microscopy (TEM) visualized long fibrils at pH 5.0 but not at pH 7.5, and these pre-formed fibrils dissolved after incubation at pH 7.5, confirming ARA1 reversibility (Fig. 2F). Finally, we employed small-angle X-ray scattering (SAXS) to characterize the solution structure of the ARA1 peptide at different pH (Fig. 2G, Table S6). The recorded spectrum at pH 7.5 was best described by a random coil model, which implies flexible, unfolded chains in solution. Fitting these data to a Gaussian coil model (Debye, 1947; Glinka, 2001) yielded a radius of gyration (R_g_) of 7.8 Å. This value is consistent with the theoretical prediction for a 15-residue peptide, supporting the conclusion that at pH 7.5 ARA1 is in a monomeric, non-aggregated state. In contrast, a striking change in the scattering profile was observed at pH 5.0, reflecting a regular cylindrical structure with an elliptical cross-section (Pedersen and Schurtenberger, 1996; Chen, Butler and Magid, 2006) (Fig. 2G, Table S6), similar to other amyloid fibrils like Aβ42 (Lattanzi *et al*., 2021). Additional parameters, including the minor radius of 29.4 Å, the axis ratio of 3.6 for the elliptical cross-section, and the clear form factor oscillation peak (Table S6), support the conclusion that the ARA1 peptide forms highly regular elongated structure with sharp solvent-amyloid interfaces. Taken together, we conclude that ARA1 responds to pH changes by protonation of its central histidine, forming reversible amyloid fibrils at low pH, while staying monomeric at pH 7.5.

We designed a series of ARA1 mutants to validate the importance of the residues surrounding the central histidine and explore the contribution of other protonatable residues (D/E) (Fig. 2H, Table S1). As predicted, ThT binding assays confirmed that the highest scoring peptide (ARA^const1^) with 3 negatively charged and 2 neutral residues mutated to hydrophobic amino acids, converts the reversible ARA1 peptide into a constitutive amyloid. More subtly, mutating one or multiple negatively charged aspartate and glutamate residues to neutral asparagine or glutamine, respectively, resulted in a poorly reversible amyloid (ARA^irrev^) or a constitutive amyloid (ARA^const2^). Conversely, ThT staining was lost as predicted with the lowest scoring peptide (ARA^non^), in which two hydrophobic residues were mutated to glycine (Fig. 2I). Therefore, this mutational analysis validates the central histidine as the key pH-sensor, while the surrounding negatively charged and hydrophobic amino acids fine-tune the level of pH-sensing and reversibility.

### Yeast proteins with predicted pH-sensing amyloid cores assemble in an SDS-resistant and pH-dependent manner during stationary phase

To explore reversible amyloids *in vivo*, we applied the bioinformatic workflow to the proteome of budding yeast *Saccharomyces cerevisiae*, which revealed 633 pH-sensing amyloid core peptides in 576 proteins (Fig. 3A, Table S4). Many proteins with predicted amyloid cores were homologous between yeast (188 proteins, 33%) and humans (Fig. 3B). The GO term analysis revealed enrichment of DNA-associated processes in both organisms whereas human candidates were particularly enriched in ATP-binding proteins and components regulating microtubule dynamics (Fig. S3A-B, Table S7).

**Figure 3:**
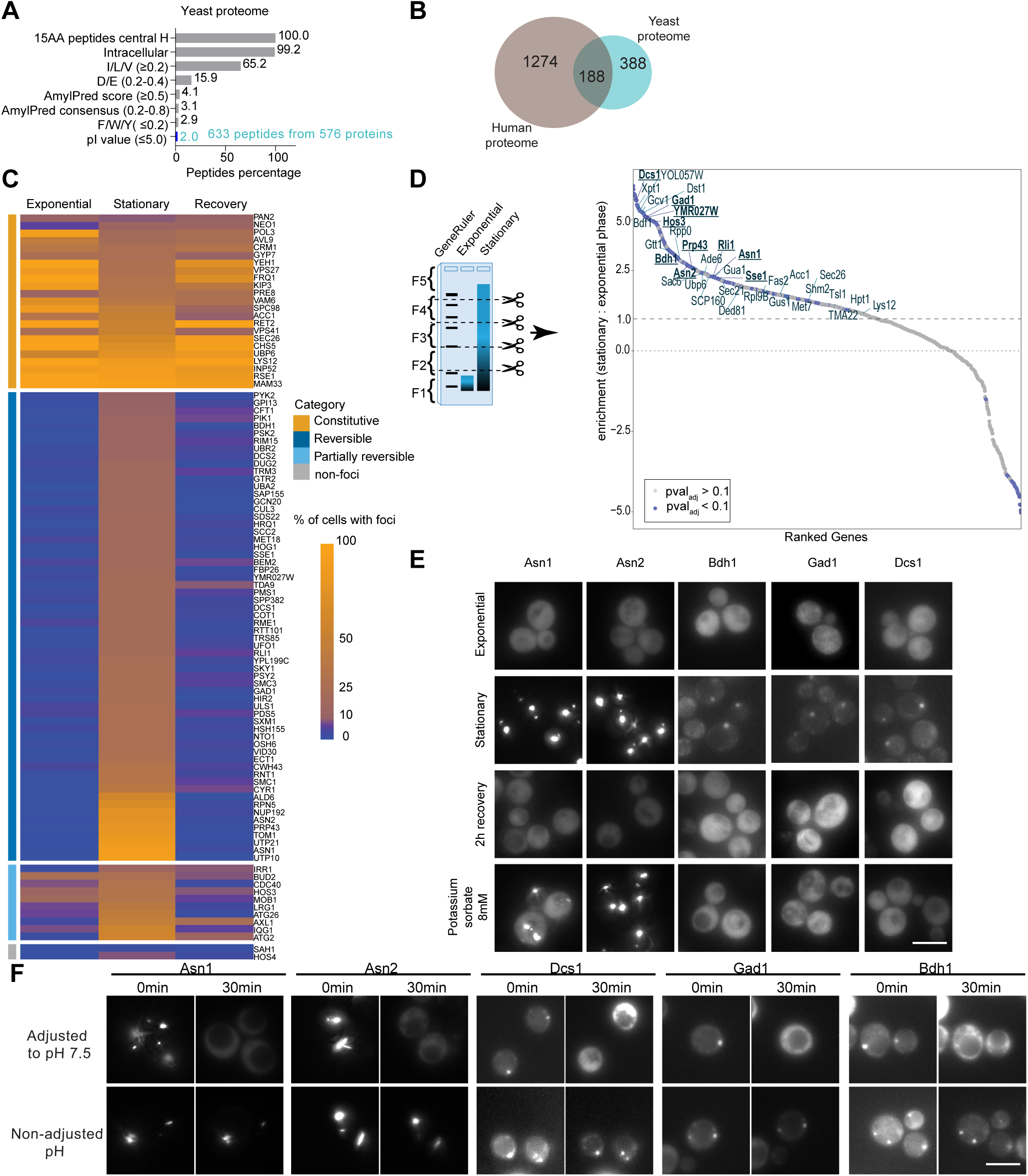
Yeast proteins with predicted pH-sensing amyloid cores assemble in an SDS-resistant and pH-dependent manner during stationary phase. A. The bioinformatic workflow (Fig. 1E) was applied to the yeast proteome, and total 15-mer peptides with a central histidine and no prolines is represented as 100%. The percentage (%) of retained peptides after sequentially applying the selection criteria is plotted, resulting in 633 peptides (2%) among 576 proteins that qualify as pH-sensing amyloid core candidates. B. Venn diagram illustrates the comparative bioinformatic analysis of the predicted yeast (576 proteins) and human (1’523 proteins) datasets. The overlapping subset contains 188 yeast proteins corresponding to 249 human homologous proteins that contain the predicted pH-sensing amyloid cores in both. C. Heat map of 97 endogenously GFP-tagged yeast proteins visualized by microscopy during exponential growth, after 2 days in stationary phase, or 1 h after recovery from 2-day stationary phase in full media. At least 100 cells were quantified per each condition in two independent replicates. Proteins were categorized as constitutive, reversible, partially reversible and non-foci based on the percentage (%) of cells with visible foci: constitutive: stationary >10%, log >5%, recovery >10%; reversible: stationary >10%, log <5%, recovery <10%; non-foci: stationary <10%, log <5%, recovery <10%; partially reversible: remaining proteins with reduced foci after recovery compared to stationary phase. D. Cartoon illustrating the fractionation of SDD–AGE gels (left). Yeast extracts from exponentially growing and 6-day stationary phase cells were separated on SDS-PAGE containing 1% SDS. Gel slices were identified by the GeneRuler (GR) molecular weight marker and subjected to MS analysis. Protein abundance (right) is calculated as the enrichment (E) in the high-molecular weight (5^th^) fraction in stationary (stat) compared to exponential (exp) phase [E= log_2_((5^th^ fraction_stat_/all fractions_stat_) : (5^th^ fraction_exp_/all fractions_exp_))]; n=3, ROTS, FDR = pval_adj_≤0.1 as significance threshold. E. Representative fluorescence microscopy images of cells expressing mNG-tagged Asn1, Asn2, Bdh1, Gad1 or Dcs1 during exponential growth in full media, after 2 days in stationary phase and following 2 h recovery from stationary phase cells in full media. Where indicated, the intracellular pH was artificially decreased in exponentially growing cells by adding 8 mM potassium sorbate for 30 min; n=3, scale bar: 5 µm. F. Representative fluorescence microscopy images of 3-day stationary phase cells expressing mNG-tagged Asn1, Asn2, Bdh1, Gad1 and Dcs1 exposed to stationary phase media. The medium was then adjusted (pH 7.5) to increase intracellular pH or non-adjusted for control. The cells were imaged at time 0 or after 30 min, when the intracellular pH equilibrated to the pH of the extracellular medium; n=3, scale bar: 5 µm. Note that elevating the intracellular pH of stationary phase cells is sufficient to dissolve foci.

To test whether the predicted core motifs form reversible assemblies *in vivo* in the full-length context, we carried out two complementary assays. First, using fluorescence microscopy, we screened 97 conserved yeast candidates using the Euroscarf GFP-fusion strain collection (Fig. 3C, Table S8) by observing foci formation in stationary phase cells, as the intracellular pH is known to decrease upon energy depletion conditions (Imai and Ohno, 1995; Dechant *et al*., 2010; Munder *et al*., 2016), compared to exponentially growing cells and recovery phase, when pH-dependent foci are not expected (Cereghetti *et al*., 2024). Indeed, almost 75% of these GFP-tagged proteins formed cytoplasmic foci after 2 days in stationary phase (Fig 3C). Most foci were reversible upon addition of fresh media, with a subset partially reversible. 23 proteins formed constitutive foci and two failed to assemble. Second, we used a global approach to screen for proteins that form SDS-resistant structures upon starvation (Fig. 3D). Extracts from either exponentially growing or 6-day stationary phase cells were separated by semi-denaturing detergent agarose gel electrophoresis (SDD-AGE), and proteins extracted from gel slices were quantified by MS analysis (Fig. 3D, S3C). This assay detected 551 proteins that significantly shifted into the highest molecular weight fraction in stationary phase (Fig. 3D, Table S9). These proteins are annotated with GO terms such as “stress granule (SG)”, “protein translation” or “amino acid biosynthetic process” (Fig. S3D, Table S7). These results showed significant overlap with previously published literature on aggregation-prone proteins, including insoluble proteomes detected by SDS-resistance assay in aged yeast (Peters *et al*., 2012), in cells at mid-exponential phase (Chen *et al*., 2021), in proteins pelleted in stationary phase (Narayanaswamy *et al*., 2009) or forming foci upon artificial intracellular pH decrease (Munder *et al*., 2016), but also with SG proteins (Jain *et al*., 2016) or proteins pelleted upon acute heat shock (Cherkasov *et al*., 2015) (Fig S3E, Table S9). In contrast, no enrichment for prion domains or P-body proteins was detected (Alberti *et al*., 2009; Jain *et al*., 2016). Among the SDD-AGE hits, 35 proteins contained a predicted pH-sensing amyloid core motif, and most (10) from the ones included in our microscopy screen were confirmed to form starvation-induced, reversible foci (Fig. 3C and D). Of these 10, we selected a few for additional analysis, including the metabolic enzymes ATP-dependent asparagine synthetases Asn1 and Asn2, the butanediol dehydrogenase Bdh1, the glutamate decarboxylase Gad1, and the mRNA decapping scavenger enzyme Dcs1. Asn1 and Asn2 share 88% sequence identity and synthesize the non-essential amino acid asparagine, and are known to form cytoophidia upon starvation (Zhang *et al*., 2018; Noree, Sirinonthanawech and Wilhelm, 2019). Indeed, these mNeonGreen (mNG)-tagged candidates formed foci in stationary phase cells, which were rapidly resolved upon refeeding (Fig. 3E, S3F). Moreover, these foci rapidly dissolved when shifting the acidic pH of the stationary phase media to a near-neutral value, thus causing an increase in intracellular pH (Fig 3F, S3H), implying that a low intracellular pH is necessary to maintain visible foci. Interestingly, Asn1 and Asn2, but not Gad1, Bdh1 and Dcs1 rapidly formed foci upon artificially lowering intracellular pH using potassium sorbate (Fig. 3E, S3G), suggesting that low intracellular pH is sufficient to trigger the formation of Asn1 and Asn2 foci, whereas additional signals are needed for Gad1, Bdh1 and Dcs1, possibly to expose the amyloid core motif. Taken together, these data demonstrate that these metabolic enzymes reversibly assemble into stable structures in a pH-dependent manner upon stress.

### Asn1 fibrillation requires pH-dependent protonation of the central histidine in the amyloid core motif

We focused biochemical and biophysical follow up on Asn1 (Fig. 4A), as low pH is sufficient to trigger its reversible assembly *in vivo* (Fig. 3E). As predicted, the Asn1_core_^WT^ peptide forms fibrils at low pH, due to the protonation of its central histidine with a pKa value of 6.49±0.16 (Fig. S4A-B). The corresponding NMR signal intensity of the soluble fraction decreases as the peptide becomes insoluble at lower pH (Fig. 4SC). Moreover, SAXS measurements of the Asn1_core_^WT^ peptide at pH 5.0 displayed a scattering profile best modelled as a flexible cylinder (Pedersen and Schurtenberger, 1996; Chen, Butler and Magid, 2006), while at pH 7.5 non-aggregated monomers were observed, according to the Gaussian coil model (Fig. S4D, Table S6). CD measurements (Fig. S4E), TEM-imaging (Fig. S4F) and ThT binding assay (Fig. S4G, Table S1) further confirmed that the Asn1_core_^WT^ peptide forms pH-sensing reversible amyloid fibrils. As expected, the designed mutants, Asn1_core_^non^, with histidine mutated to non-protonatable alanine, and Asn1_core_^agg^, with negatively charged D/E residues mutated to the corresponding neutral N/Q residues (Fig. 4A), prevented or promoted the formation of amyloid structures, respectively (Fig. S4F-G, Table S1).

**Figure 4:**
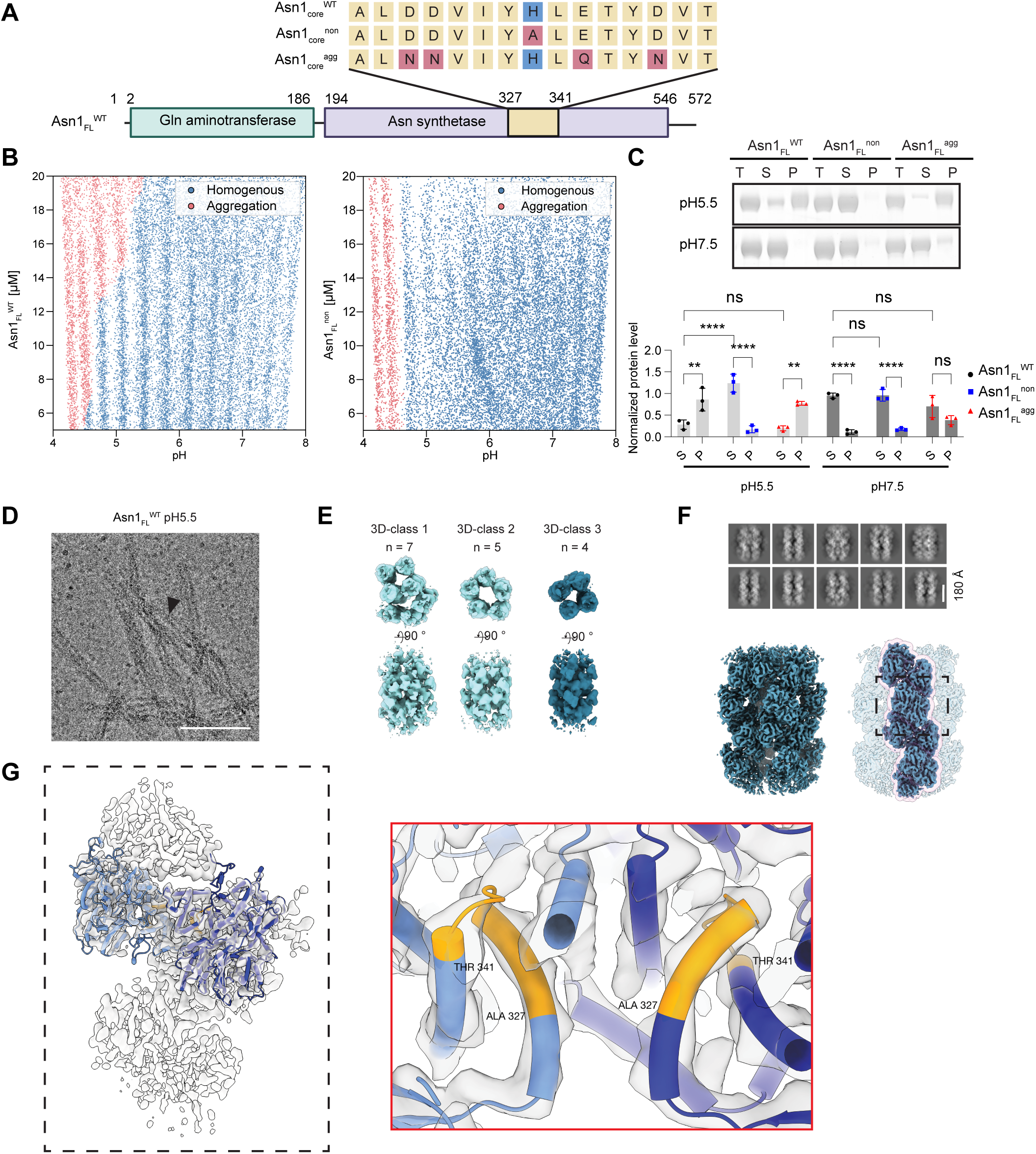
Protonation of the central histidine in the amyloid core promotes full length Asn1 fibril formation. A. Schematic representation of full length (FL) Asn1, highlighting the predicted amyloid core motif (inset). Mutating the central histidine (H334, blue) to a non-protonatable alanine is expected to interfere with fibrillation (Asn1_core_^non^), while mutating the negatively charged residues to the corresponding neutral amino acids (D329N, D330N, E336Q and D339N) is predicted to generate a more aggregation-prone mutant (Asn1_core_^agg^). The mutations are highlighted in red, and the corresponding core peptides were characterized in Fig. S4. B. High resolution phase diagrams of Asn1_FL_^WT^ or Asn1_FL_^non^ titrated at pH range 4-8. Purified full-length proteins were labelled with Alexa Fluor 647, encapsulated as water-in-oil microdroplets at the indicated pH and imaged. Each datapoint represents an individual microdroplet classified as either homogeneous (blue) or non-homogenous aggregated (red); n=19529 (Asn1_FL_^WT^), n=17981 (Asn1_FL_^non^) microdroplets. C. Purified Asn1_FL_^WT^, Asn1_FL_^non^ or Asn1_FL_^agg^ proteins were incubated for 24 h at 25°C at pH 5.5 or pH 7.5 and then separated by centrifugation into supernatant (S) and pellet (P) fractions. A sample of the total (T) protein before centrifugation was included. Proteins were visualized by Coomassie blue staining (representative image, upper panel), and the mean grey value of S and P was normalized by respective T to quantify the relative protein levels for each condition (bottom panel); n=3, statistical significance assessed by a two-way ANOVA test, followed by Tukey’s multiple-comparisons, ** p <0.01, *** p <0.001, **** p <0.0001; ns, not significant. D. A representative cryo-EM micrograph showing Asn1 fibril bundles after incubating purified Asn1_FL_^WT^ for 24 h at 25°C at pH 5.5. Single protofilaments assembling into bundles are indicated by the black arrow; scale bar: 150 nm. E. Asn1_FL_^WT^ assembles polymorphic fibrils divided in three distinct classes, composed of 7 (3D-class 1), 5 (3D-class 2) or 4 protofilaments (3D-class 3). F. Representative 2D class averages of Asn1_FL_^WT^ fibrils from 3D-class 3 (top images). Note that Asn1 fibrils do not adopt a conventional cross-β-sheet structure characteristic for classical amyloids. The cryo-EM density map of a 3D-class 3 Asn1_FL_^WT^ fibril (bottom) displayed as complete fibril (left) and with a single protofilament highlighted (right). The dashed section of the protofilament was used to generate a mask for focused refinement and model building shown in panel G. G. Refined model of a Asn1_FL_^WT^ dimer was fit to the 4.8 Å cryo-EM map of a single protofilament (PDB ID: 31ZZ). The model reveals tight stacking of Asn1 monomers along the protofilament axis. Residues 327–341 within the dimer interface corresponding to the amyloid core motif are highlighted in gold (inset, red box).

To corroborate these data, we next purified full length (FL) recombinant Asn1_FL_^WT^, Asn1_FL_^non^ and Asn1_FL_^agg^ from *E. coli*. Indeed, Asn1_FL_^WT^ rapidly aggregated in a concentration-and pH-dependent manner, starting the transition around pH 5.5, as determined in a microfluidic oil encapsulation system (Arter *et al*., 2022; Agarwal *et al*., 2025) (Fig. 4B). In contrast, Asn1_FL_^non^ did not show a concentration-dependent transition, and aggregates were observed only at non-physiological pH values below 4.5. Consistent with this analysis, purified Asn1_FL_^WT^ was fully soluble at pH 7.5, but readily pelleted at low pH (Fig. 4C). In contrast, purified Asn1_FL_^non^ remained soluble at pH 5.5, whereas Asn1_FL_^agg^ already partially pelleted at pH 7.5 and completely aggregated at pH 5.5. Together these results show that Asn1_FL_^WT^ aggregates in a pH-dependent manner, and the protonatable histidine residue in its amyloid core facilitates this transition.

To examine the molecular structure of pH-sensing Asn1 aggregates, purified Asn1_FL_^WT^ protein was pre-incubated at pH 5.5 and analyzed with single-particle cryo-EM (Fig. 4D–G, Fig. S5). Single-particle analysis (SPA) revealed multiple fibril morphologies, which could be grouped into three distinct 3D-classes based on the number of protofilaments (Fig. 4D-E). These included bundled fibrils composed of either seven (3D-class 1) or five protofilaments (3D-class 2), as well as single fibril consisting of four protofilaments (3D-class 3). The higher-order fibril assemblies observed in 3D-class 1 and 3D-class 2 resemble the bundling behavior described for yeast CTP synthases (Ura7 and Ura8), which was proposed to inactivate their enzymatic activity under starvation (Hansen *et al*., 2021). We focused our analysis on a single protofilament from the 3D-class 3 and obtained a 3D-reconstruction reaching an overall resolution of 4.8 Å (Fig. 4G), which allowed building an atomic model (PDB ID: 31ZZ) based on an AlphaFold3-predicted Asn1_FL_^WT^ structure. This model, together with reconstructions of higher-order protofilament bundles, revealed that Asn1_FL_^WT^ assembles into fibrils with a regular repeat of two monomers along the fibril axis (Fig. 4F), and thus does not resemble a classical cross-β-sheet amyloid fibril architecture. The amyloid core motif (A327-T341) does not appear to stack directly onto the same motif in adjacent subunits, suggesting that the structural arrangement in the context of the full-length protein differs from that observed in isolated core peptides (Fig. 4G). Instead, the Asn1_FL_^WT^ fibril model indicates that the amyloid core region encompassing the protonatable H334 residue directly regulates monomer–monomer interactions. Together, these structural features provide a molecular framework for understanding how pH-dependent histidine protonation of the identified amyloid core motif modulates Asn1 fibril assembly and disassembly.

### Regulated assembly of Asn1 allows timely growth restart from stationary phase

To understand the physiological function of pH-dependent reversible Asn1 fibrils in yeast, we generated yeast strains harboring the SG marker Ded1-mCherry (Hilliker *et al*., 2011) and endogenously expressing mNG-tagged wild-type Asn1 (Asn1^WT^), or its non-aggregating (Asn1^non^) or aggregation-prone (Asn1^agg^) mutants (Fig. 4A). As expected, Asn1^WT^ and Asn1^non^ were dispersed in the cytoplasm in exponentially growing cells (Fig. S6A), while after 3 days in stationary phase only Asn1^WT^ assembled elongated foci previously described as cytoophidia (Zhang *et al*., 2018), which did not co-localize with Ded1-positive SGs (Fig. 5A). The number of cells with visible Asn1^WT^ foci was reduced when cells were grown into stationary phase in minimal media, containing only essential amino acids (Fig. S6B), suggesting that metabolite availability may affect the assembly of Asn1^WT^ cytoophidia. Importantly, Asn1^non^ failed to form visible foci in stationary phase (Fig. 5A), demonstrating that starvation-induced protonation of the amyloid core motif is required to trigger reversible fibrillation *in vivo*. Surprisingly, significantly less cells with Ded1 foci were observed in Asn1^non^ strain in the complete but not the minimal media (Fig. 5A, S6B), indicating crosstalk between SG formation and Asn1 cytoophidia during stationary phase. Asn1^agg^ was mostly dispersed in exponentially growing cells (Fig. S6A) but formed aberrant structures co-localizing with SGs in stationary phase (Fig. 5A). Interestingly, Asn1^WT^ but not Asn1^non^ rapidly formed foci when exponentially growing cells were treated with potassium sorbate (Fig. 5B, S6C), implying that lowering pH is sufficient to trigger assembly *in vivo*. Asn1^agg^ showed substantial foci formation already at lower potassium sorbate concentrations, indicating that it is more aggregation prone than Asn1^WT^. Conversely, low pH was necessary to maintain the Asn1^WT^ cytoophidia formed in stationary phase, as adjusting the extracellular pH to a near-neutral value, and as a result elevating the intracellular pH, led to foci dissolution (Fig. S6D). Together, we conclude that Asn1 requires the protonatable histidine 334 in its amyloid core motif to assemble in a pH-dependent manner during stationary phase, and lowering pH is necessary and sufficient for foci formation *in vivo*.

**Figure 5:**
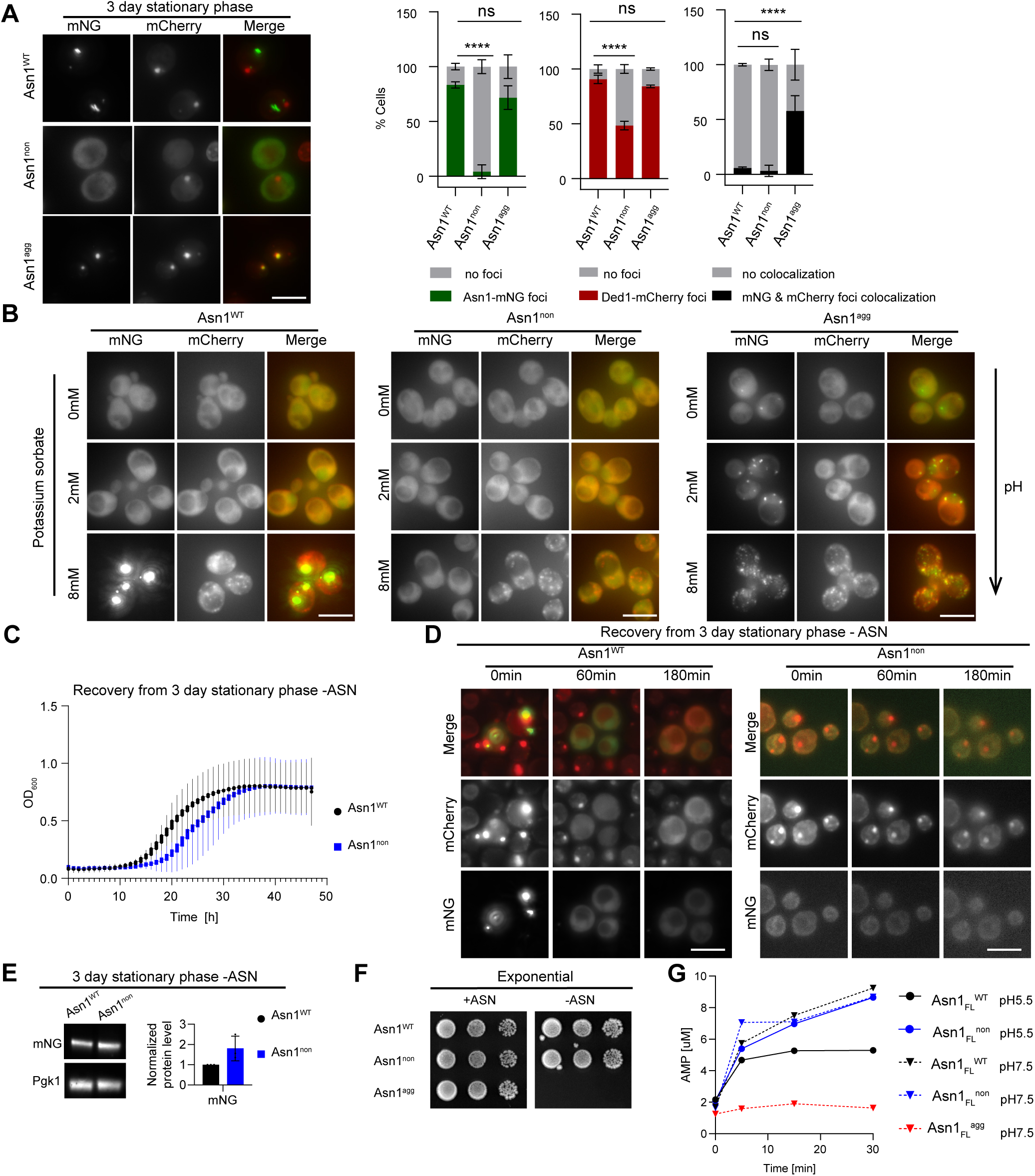
Regulated pH-dependent Asn1 assembly allows cells to efficiently restart growth after release from stationary phase. A. *asn1Δ* cells harboring the SG marker Ded1-mCherry and stably expressing mNG-tagged Asn1^WT^, Asn1^non^ or Asn1^agg^ from the endogenous promoter were imaged at 3-day stationary phase in complete media (representative images, left). The bar plots (right) depict the percentage (%) of cells with mNG and mCherry foci, and their co-localization; n=3, mean ± SD. A two-way ANOVA followed by Dunnett’s multiple-comparisons test controls statistical significance. **** p <0.0001 vs. WT; ns, not significant, scale bar: 5 µm. B. Exponentially growing *asn1Δ* cells harboring Ded1-mCherry and stably expressing mNG-tagged Asn1^WT^, Asn1^non^ or Asn1^agg^ from the endogenous promoter in complete medium were treated with 0, 2 or 8 mM potassium sorbate for 30 min to lower intracellular pH. Fluorescence microscopy was performed to visualize foci formation of Asn1 (mNG) and Ded1 (mCherry), and their colocalization was analyzed in the merge; representative images, n=3; scale bar: 5 µm. Note that with equivalent exposure the Asn1^WT^ foci are saturated. Quantification is shown in Fig. S6C. C. After 3-day stationary phase in minimal medium without asparagine (-ASN) *asn1Δ* cells stably expressing Asn1^WT^ or Asn1^non^ from the endogenous promoter were diluted into fresh medium (-ASN), and the recovery was monitored by optical density (OD_600_) measurements. Note that *asn1^non^*cells exhibit delayed growth restart compared to *asn1^WT^* controls, despite the presence of Asn2; n=3, mean ± SD. D. The recovery of 3-day stationary phase *asn1Δ* cells harboring Ded1-mCherry and stably expressing mNG-tagged Asn1^WT^ or Asn1^non^ from the endogenous promoter in fresh SD-medium without asparagine (-ASN) was monitored by microscopy at the times indicated. Note that disassembly of SGs marked by Ded1-mCherry was delayed in *asn1^non^* cells; representative images, n=3, scale bar: 5 µm. Quantification is shown in Fig. S6F. E. Extracts of *asn1Δ* cells harboring Ded1-mCherry and stably expressing mNG-tagged Asn1^WT^ or Asn1^non^ from the endogenous promoter were prepared after 3 days stationary phase in minimal media without asparagine (-ASN). The levels of mNG-tagged Asn1^WT^ or Asn1^non^ proteins were monitored by immunoblotting with mNG-specific antibody (representative blot, left panel). Pgk1 controlled equal loading. The band intensity was quantified and normalized to Pgk1 levels (right panel); n=3, mean ± SD. F. Representative serial dilution spottings of exponentially growing *asn1Δ asn2Δ* double mutant cells stably expressing Asn1^WT^, Asn1^non^ or Asn1^agg^ on minimal media with (+ASN) or without (-ASN) addition of asparagine; n=3. Plates were photographed after 3 days at 30°C. G. *In vitro* enzymatic activity assays with purified Asn1_FL_^WT^, Asn1_FL_^non^ or Asn1_Fl_^agg^ protein, preincubated for 24 h at 25°C at pH 5.5 or pH 7.5. After addition of ATP at time 0, the production of AMP was measured every 10 min at pH 7.5 and/or pH 5.5; n=2, mean ± SD.

To investigate the functional relevance of Asn1 assembly and SG dynamics, we released *asn1^WT^* and *asn1^non^* strains after 3 days in stationary phase into fresh minimal media (Fig. 5C). Importantly, OD_600_ measurements showed that cells unable to form Asn1 foci showed delayed growth recovery compared to wild-type controls (Fig. 5C), despite the presence of wild-type Asn2. No such recovery delay was observed in *asn1Δ* strains (Fig. S6E). Moreover, in these conditions, while in wild-type cells both Ded1 and Asn1 foci rapidly dissolve, Ded1 foci persist significantly longer in *asn1^non^* cells, indicating that SG disassembly is altered with an Asn1 assembly defect (Fig. 5D, S6F). Importantly, immunoblotting revealed that Asn1^non^ is not degraded during stationary phase (Fig. 5E). Both the delayed growth recovery and the delayed Ded1 foci disassembly in *asn1^non^* cells can be rescued by the addition of asparagine to the media (Fig. S6G-I). However, spotting assays and growth analysis of *asn1Δ asn2Δ* double mutants expressing Asn1^non^ or Asn1^WT^ showed growth in minimal media without asparagine (Fig. 5F, S6J), implying that the Asn1^non^ mutant protein is catalytically active *in vivo*, while the *asn1Δ asn2Δ* double deletion alone is not viable (Fig. S6K). Importantly, *in vitro* assays confirmed that the catalytic activity of purified Asn1^non^ enzyme is comparable to Asn1^WT^ when measured at pH 7.5 (Fig. 5G). However, Asn1^non^ activity remains high when the pH is dropped to 5.5, in contrast to Asn1^WT^ enzyme which forms fibrils under these conditions (Fig. 5G). We conclude that Asn1 fibrils are catalytically inactive, while the non-aggregating Asn1 mutant fails to assemble fibrils and thus remains active during low pH conditions such as stationary phase. In contrast, the Asn1^agg^ mutant is catalytically inactive *in vitro* at pH 7.5 (Fig. 5G) and it cannot sustain growth of *asn1Δ asn2Δ* strains without addition of asparagine in the media (Fig. 5F, S6K). Taken together, these results suggest that the formation of Asn1 cytoophidia inhibits Asn1 activity during stationary phase, and this mechanism is functionally relevant to allow for a timely growth restart after stress release.

## Discussion

### Bioinformatic and experimental workflow to identify regulatable amyloid peptides

Taking advantage of biophysical properties beyond sequence conservation, and experimental validation of supplemental criteria using the TAAPAS pipeline, allowed us to establish a computational workflow that accurately predicts pH-sensing amyloid core motifs. This approach is adaptable and scalable, and amenable to increase throughput by partial automation, thus offering as-of-yet unrealized potential in amyloid research. For example, the pipeline can be adapted to (1) identify reversible amyloid cores throughout the tree of life to evaluate conservation and the evolution of specific motifs, (2) probe amyloid assembly and disassembly kinetics, (3) test conditions beyond differential pH, for example post-translational modifications, metal ions or salt concentrations, or (4) assess additional readouts like further dye-binding assays, microscopy or catalytic activity measurements. Mining the TAPAAS peptide library may also provide detailed information about underlying assembly and disassembly mechanisms, like different pH-value set points or hysteresis, as recently demonstrated for the PKM2 core (Dear *et al*., 2025). Adjustment to the experimental set up may also reduce technical caveats, including limited resolution of the MS readout or possible ThT off-target effects (Biancalana and Koide, 2010). Thus, TAAPAS is suitable as a powerful screening platform for biotech or medical applications.

### The molecular grammar and regulation of amyloids

Our data provides evidence that pH-sensing can be a more wide-spread regulatory mechanism than previously thought, in line with earlier reported pH-sensing amyloids (Maji *et al*., 2009; Morris *et al*., 2023; Cereghetti *et al*., 2024). We have characterized the molecular properties of amyloid core motifs with a central histidine that when protonated acts as a molecular switch for reversible assembly. In a polypeptide context, the average pKa of histidine is 6.6 (Grimsley, Scholtz and Pace, 2009), consistent with our measurements on amyloid core peptides and within the physiological pH fluctuation range. While constitutive amyloid cores mainly use hydrophobic amino acids, a distinguishing feature of pH-regulated cores is the overall high content of acidic side chains, supporting the view that acidic residues act as gatekeepers for protein aggregation (Houben *et al*., 2020; Horváth *et al*., 2023). Precisely how these glutamate and aspartate residues affect amyloid fibril formation remains unclear, and moreover, until now we do not know if they are protonated in fibrils. If unprotonated, the electrostatic repulsion might prevent amyloid assembly until the pH-sensing histidine becomes protonated. Unprotonated glutamate or aspartate may also stabilize assembly at low pH by interacting with the protonated histidine. Alternatively, if glutamate or aspartate become protonated, they may form a “polar zipper” themselves as predicted for β-synuclein (Moriarty *et al*., 2017), and/or electrostatic repulsion may diminish, as proposed for the PKM2 core (Dear *et al*., 2025). Although pKa values of glutamate and aspartate in isolation are low (∼4.3 and 3.9, respectively), they can increase substantially within proteins (Grimsley, Scholtz and Pace, 2009). Accordingly, the pKa of glutamate buried in β-endorphin fibrils was predicted to be above 5.5, and simulations indicated progressive deprotonation at neutral pH and resulting electrostatic repulsion mediated fibril disassembly (Liu *et al*., 2021). Higher physiological pKa values could enable protonation, for example during stress-induced intracellular acidification in yeast (Dechant *et al*., 2010). Altogether, the available data support the view that the protonation state of titratable residues within the amyloid core motifs serves as a molecular switch controlling amyloid assembly and disassembly. Such a protonation-dependent mechanism is passive and possibly evolutionary ancient (Tze-Fei Wong, 1975).

Other factors beyond protonation can regulate fibril formation. The electrostatic landscape of a polypeptide can be reshaped for example by phosphorylation, ATP- or metal-ion binding, which were also proposed to regulate amyloids (Dong *et al*., 2007; Cereghetti *et al*., 2021; Yamaguchi *et al*., 2026), but systematic exploration remains elusive. Regulatory conformational changes and/or complex rearrangements may also be required to expose critical motifs. For example, the amyloid core of Cdc19 maps to its tetramer interface, and monomerization triggered by decreased FBP levels precedes fibril assembly (Saad *et al*., 2017; Cereghetti *et al*., 2021). Indeed, while for Asn1 lowering the cytosolic pH was sufficient to trigger fibril formation, other candidates required additional starvation signals to assemble foci *in vivo.* Although Asn1 and Asn2 both possess identical amyloid cores, Asn1 is required to assemble starvation-induced Asn2 foci (Noree, Sirinonthanawech and Wilhelm, 2019). It is possible that *in vivo* the amyloid core motif of Asn2 is not accessible in the absence of Asn1, or Asn1 may act as structural scaffold, allowing Asn2 to assemble into heterogenous Asn1/2 fibrils. Taken together, the physiological context likely determines the functional response of pH-sensing amyloid core motifs.

### pH-sensing amyloid core motifs may regulate diverse assemblies

We found multiple yeast proteins with the predicted pH-sensing amyloid cores that formed foci and/or SDS-resistant assemblies during stationary phase, when pH decreases (Imai and Ohno, 1995; Joyner *et al*., 2016; Munder *et al*., 2016). Our analysis revealed structural distinctions from the prototypical cross-β-sheet fold in the full-length context, coherent with the notion that although pathological amyloids are notoriously stable, functional amyloids reveal surprising diversity (Fowler *et al*., 2007; Pham, Kwan and Sunde, 2014; Otzen and Riek, 2019). Full-length Asn1 fibrils are composed of unbranched protofilaments formed by regular repeats of two monomers, reminiscent of other filamentous proteins forming cytoophidia such as inosine monophosphate dehydrogenase (IMD) (Johnson and Kollman, 2020) or CTP synthase (CTPS) (Hansen *et al*., 2021). Interestingly, protonation of the histidine in the Asn1 amyloid core motif directly regulates monomer–monomer interactions but does not cause stacking of cross-β-sheets, classical for amyloids. Still, full-length Asn1 fibrils exhibit amyloid-like characteristics, including aggregation (Fig. 4) and SDS-resistance reporting on fibril stability (Fig. 3). Thus, rather than a generic amyloid fold, pH-sensing amyloid core motifs may exploit physical principles to assemble an array of fibrillar structures with diverse architecture influenced by apolar, polar, ionic and hydrogen bonding interactions, together with co-factor binding and locally disordered regions. Clearly further structural studies are warranted to reveal the conformational diversity and function of amyloid core motifs in different species. Finally, as discovery progresses the amyloid nomenclature may need reevaluation to include the expanding structural diversity and polymorphism.

### pH-dependent regulation of Asn1 fibrils is required for efficient recovery from stationary phase

Recent work revealed that some metabolic enzymes polymerize into reversible fibrils, constituting a conserved mechanism to compartmentalize and regulate biochemical reactions (Park and Horton, 2019). For example, aldehyde dehydrogenase Ald4 assembles mitochondrial filaments, while acetyl-coenzyme A synthetase Acs1 forms cytoplasmic fibrils that bind ADP and facilitates cell cycle re-entry of starved cells (Hugener *et al*., 2024). While different signals trigger filament assembly, stress-induced lowering of the intracellular pH regulates the assembly of reversible cytoophidia, including Asn1, CTPS and glutamine synthetase Gln1 (Petrovska *et al*., 2014; Noree *et al*., 2019; Hansen *et al*., 2021). Polymerization of CTPS inhibits its catalytic activity in yeast (Hansen *et al*., 2021), and similarly we found that Asn1 fibrils are catalytically inactive, thus likely blocking asparagine production during stationary phase. Importantly, cells defective for Asn1 fibril formation exhibit recovery problems after refeeding (Fig.5). However, Asn1^non^ is not degraded during stationary phase, indicating that these cells may fail to fully inactivate Asn1. This recovery delay is rescued by addition of asparagine in the culture medium, implying that asparagine or a metabolite dependent on the presence of asparagine becomes limiting if Asn1 remains active during starvation. Surprisingly, this defect also delays SG disassembly, indicating crosstalk among metabolic pathways during stationary phase. Finally, mutating the Asn1 amyloid core motif to mimic constitutive fibril formation rendered the full-length Asn1 mutant protein catalytically inactive and the misfolded protein accumulated in SGs, likely to prevent cell toxicity. Further work will be required to understand how aggregation-prone Asn1 assemblies are recognized and targeted to SGs during starvation, and how these aggregates are cleared after stress release and SG disassembly. Irrespective, our work uncovered a widespread mechanism that regulates reversible protein assembly and quality control during starvation and possibly other stress conditions.

## Author contributions

Conceptualization: A.K., C.S., D.P. and M.P.; funding acquisition: A.K., N.D., T.P.J.K. and M.P.; bioinformatics: A.K., D.P., G.C. and I.K.; TAPAAS: D.P. and F.U.; peptide analysis: assays - A.K. and D.P., SAXS - A.K. and V.L.B., CD - S.K., NMR - A.G.; SDD-AGE: experimental design - A.K., C.S., D.P. and F.U., assay and proteomics - A.K. and F.U., data analysis - A.K., D.P. and F.U.; *in vitro* protein analysis: purification - A.K., C.S. and S.K., biochemistry - A.K.; cryo-EM analysis: M.B., P.A., and T.C.; microfluidics: experimental design - G.C and T.A., protein labelling - G.C., assay and data analysis - T.A.; yeast analysis: strain construction - A.K. and C.W., strain characterization and microscopy - A.K., C.W. and D.P., with ScopeM; data visualization: A.K., C.W. and D.P, with inputs from all authors; supervision: N.D., T.P.J.K. and M.P.; writing: original draft - A.K., D.P., and M.P., review and editing - A.K., C.W., D.P. and M.P., with inputs from all authors.

## Acknowledgements

We thank the ETH Scientific Center for Optical and Electron Microscopy (ScopeM) and in particular Miroslav Peterek for assistance with ns-TEM, Sung Sik Lee for help with fluorescence microscopy, Davide Amendola and Julius Rabl for advising on cryoEM sample preparation and model refinement, and Michał Okoniewski from the Computational and Data Science Support for support with statistical analysis. The synchrotron SAXS data were collected with the help of Aleksi Sutinen at the P12 beamline operated by EMBL Hamburg at the PETRA III storage ring (DESY, Hamburg, Germany). This work benefited from the use of the SasView application, originally developed under NSF award DMR-0520547. SasView contains code developed with funding from the European Union’s Horizon 2020 research and innovation program under the SINE2020 project, grant agreement No 654000. We are grateful to Jinghui Luo for experimental advice, and Justin Kollman and John Calise for sharing unpublished reagents and results. We thank Gabriel Neurohr, Karsten Weis, Martin Winkler, and members of the Peter lab for helpful discussions and critical comments on the project, and Alicia Smith for editing the manuscript. F.U. acknowledges funding by a Marie Curie Individual Fellowship (Grant agreement number 101207537). T.A. and T.P.J.K. are supported by the European Research Council (ERC) [DiProPhys 101001615], and G.C. by an EMBO Fellowship (EMBO ALTF 349-2023) and the UKRI Engineering and Physical Sciences Research Council Fellowship (grant EP/Z000033/1). Work in the Peter laboratory was funded by the Swiss National Science Foundation (SNSF), the Dementia foundation, ETH Zürich and by a BIF PhD fellowship to A. K.

## Materials and Methods

### Cloning and molecular biology methods

All plasmids used in this study are listed in Table S10. Standard molecular biology methods were used.

### In silico methods

#### Bioinformatic predictions of pH- sensing amyloid cores

SLiMSearch was used to identify 15-mer peptides with central histidine and no proline residues in human and yeast proteomes, together with their respective annotations (regular expression: {7}[^P]H[^P]{7}) (Krystkowiak and Davey, 2017). Identified hits were filtered by protein localization using built-in SLiMSearch functionality. Additional filtering was performed using in-house script to apply amino acid composition criteria, calculated based on the peptide sequence, i.e. initially peptides were required to contain at least two hydrophobic and/or aromatic residues, with at least one located adjacent to the central histidine, and less than three same-charge residues clustering in a sequence. Peptides were further annotated for a low isoelectric point (pI), and amyloidogenicity using AMYLPRED2 score (ratio of positively scoring constituent methods) and consensus coverage (Tsolis *et al*., 2013). Predicted peptides were classified into three categories (pH-sensing amyloid, constitutive amyloid, non-amyloid) according to filtering criteria described in the main text. Homologous proteins were obtained from the Ensembl Compara tool and evolutionary annotations from SLiMSearch.

#### *In silico* design of artificial amyloid peptides

Position-specific scoring matrices (PSSMs) were generated from experimentally validated peptides aligned by their central histidine using the PSSMSearch tool (Krystkowiak, Manguy and Davey, 2018) (parameters: PSI-BLAST IC method, human proteome background, disorder cut-off: 0.0). For each obtained PSSM, the 5’000 highest scoring artificial peptides were generated and annotated with pI value and AMYLPRED2 scores (Tsolis *et al*., 2013). Peptides were ranked based on the PSSM p-value and those meeting predefined criteria (pI < 4.5, AMYLPRED2 consensus coverage 20-60%) were selected as the candidate synthetic peptides.

#### *In silico* design of mutant peptides

PSSMs for ARA and Asn1_core_^WT^ were created as described above. Next, based on the obtained PSSM scores, the most impacting mutations were selected, defined as a single substitution at a given position that yielded substantially lower score compared to the wild-type residue at that position. The central histidine position was excluded due to experimental constraints requiring its presence. All possible mutant variants with 1-3 mutations were generated, along with a subset of variants containing a higher number of substitutions in cases where three mutations did not break required amyloidogenic properties. Each mutated peptide was subsequently reannotated with amyloid-related attributes. Finally, the mutants were classified as constitutive, pH-sensing, or non-amyloid according to their predicted amyloidogenic properties.

#### Sequence logo generation

The sequence logos were generated with PSSMSearch tool (Krystkowiak, Manguy and Davey, 2018) using binomial log10 method with human background (disorder cut-off: 0.0.).

#### GO term enrichment analysis

GO terms enrichment analysis was performed using DAVID functional annotation tool (Huang, Sherman and Lempicki, 2009; Sherman et al., 2022). The following background lists were used: all proteins with 15-mer peptides containing central histidine, n=17’981 for human or n=5’821 for yeast predictions of pH-sensing amyloids; entire yeast proteome for SDD-AGE-MS analysis.

### *In vitro* methods with synthetic peptides

#### *In vitro* peptide samples preparation

For the initial pre-screen (Fig. S1C) lyophilized synthetic peptides with -COOH C-terminus were ordered from Shanghai GLS and dissolved to 10 mg/ml in DMSO with 10% formic acid. Peptides were stored at −20°C until further use. For the subsequent ThT binding assay, peptides were diluted in TRIS buffer (100 mM Tris-HCl, 200 mM NaCl, 1 mM MgCl_2_; pH 7.5 or 5.8) to 2.0 mg/ml, and the final pH was approximately adjusted using NaOH to ∼7.5 or low pH of ∼6.0 or less, as validated by pH indicator paper. Samples were incubated for ∼24 h at 30°C before measurements.

For all the other experiments lyophilized synthetic crude quality peptides with -CONH2 C-terminus (or -COOH terminus for experiment in Fig. S1D) were ordered from peptides&elephants GmbH, dissolved in dimethyl sulfoxide (DMSO; Sigma-Aldrich, D5879) to 10 mg/ml and stored at −20°C until further use. For all low-throughput experiments, Eppendorf Protein LoBind tubes (Eppendorf, 0030 108 442) were used. Peptides were diluted to final concentration of 0.5 mg/ml, unless stated otherwise, in sterile-filtered citrate phosphate (CPS) buffer (2 mM citric acid, 50 mM Na_2_HPO_4_, 150 mM NaCl) at the stated pH and incubated for ∼24 h at 30°C before measurements (amyloid formation tests). To test potential reversibility, samples pre-incubated for 24 h at CPS buffer pH 5.0 as described above were aliquoted (aliquots number corresponding to the number of tested pH values) and pelleted by centrifugation in RT at 21’300 rcf for 10 min. The supernatant was carefully removed, and the pellet was resuspended in an equal volume of the CPS buffer at desired pH by tapping the tube. These samples were then further incubated for 24 h at 30°C.

All peptide assays were performed in independent triplicates, unless stated otherwise.

#### ThT binding assays

##### ThT fluorescence intensity measurements

18 µl of the peptide sample were transferred in technical duplicates into a 384-well plate (Corning #3766, with nonbinding surface) and mixed with Thioflavin T (2 µl, final concentration 250 µM). ThT fluorescence intensity was measured at 30°C as single timepoint (initial experiments – Fig. S1C) or every 2 min for at least 30 min (16 timepoints) using the CLARIOstar Plus plate reader (BMG LABTECH; excitation 450 nm, emission 490 nm, 20 flashes per well, 5 s double orbital shaking at 500 rpm before each cycle).

##### Data analysis

Average or median ThT intensity value of the 16 measurement timepoints, corrected by the respective buffer with DMSO blank control, was used for data analysis. The results for the 51 initial screen peptides (Fig. S1C-D) and 25 further selected peptides for prediction validations (Fig. S1G-H) were analyzed using R software and ROTS (Reproducibility-Optimized Test Statistic, ROTS package (Suomi et al., 2017)) with default 5’000 bootstrap resampling and K equal to the number of tested peptides with false-discovery-rate (FDR) used as adjusted p value. The outcome (non-amyloid, constitutive or pH-sensing reversible or irreversible amyloid) was classified based on the arbitrary threshold of 500 a.u. (defined based on the global noise of the TAPAAS results) and significant differences in formation and/or reversibility tests at different pH. ThT binding of ARA1-3, ACA, Asn1_core_ and their mutants was analyzed in GraphPad prism 11.0.0 using statistical analysis as indicated in the figure legends. All ThT binding data was visualized using GraphPad prism 11.0.0.

#### ThT and Pelleting Assays for Amyloid Screening (TAPAAS)

##### Sample preparation

Test peptides were obtained in crude quality from peptides&elephants GmbH in a 96-well plate setup divided in 12 subsets of 46 peptides, each including repetitive PKM1, PKM2 and USP9X controls and dissolved in DMSO to 10 mg/ml. Peptides were diluted to 0.2 mg/ml (formation test) or 0.4 mg/ml (reversibility test) in CPS buffer pH 5.0 (formation and reversibility) or 7.5 (formation only) in a 96-well plate with nonbinding surface (Corning #3651). For each subset, a sample for total peptide quantification via MS was taken immediately after dissolving. For that, 3 µl of each peptide from the subset were pooled into 132 µl CPS buffer and 276 µl DMSO to obtain a final overall peptide concentration of 0.05 mg/ml. The plates were incubated for 1 day at 30°C. ThT fluorescence was measured as described above, but with single technical replicate. For the formation test the remaining peptide solution was spun down (2 h, 4’500 rcf) and the supernatant samples were taken for the MS analysis. For the reversibility test the remaining peptide solution was diluted 1:2 with CPS buffer to shift the pH to final 7.5 or remain at pH 5.0 and incubated for 1 day at 30°C under rotation at 120 rpm. Then ThT fluorescence was measured, the peptides were pelleted for 2 h at 4’500 rcf and the supernatant samples were taken for the MS analysis and ThT fluorescence from supernatant samples was measured to control for efficient pelleting. TAPAAS were performed in independent triplicate, each subdivided in respective subsets.

##### MS acquisition

All MS samples were acidified to pH below 3 immediately after collection with 30 µl of formic acid (FA, final ∼5%) and stored at −20 °C until processing. The samples were desalted using BioPureSPN Mini, PROTO 300 C18 desalting columns (Higgins Analytical Inc, USA). In brief, all centrifugation steps were performed at 50 rcf, columns were activated with 200 μl 100% acetonitrile (ACN) and washed 3x with 200 µl of 0.1% FA. Samples (420 µl ≈ 20 µq peptide material each) were loaded, and columns were washed 2x with 200 µl of 0.1% FA. Samples were eluted into fresh low-binding tubes with 300 µl of 50% ACN/0.1% FA, dried in a SpeedVac device at 45°C and stored at −20°C. Before measurements, samples were resolubilised in 10 µl of 2% ACN with 0.1% FA, vortexed for 10 s, sonicated for 10 min, and centrifuged (2 min, 21300 rcf). Supernatant was transferred to fresh low-binding tubes and 4 µl were mixed with 8 µl of master mix solution composed on 2% ACN in 0.1% FA, 0.5 pmol/µl of purified BSA and iRT standard peptides (1:50, Biognosys AG) prior to LC-MS analysis (2 µl of final 0.66 µg/µl peptide material injected).

LC-MS/MS analysis was performed on an Orbitrap QExactive+ mass spectrometer (Thermo Fisher) coupled to an EASY-nLC-1000 liquid chromatography system (Thermo Fisher). Peptides were separated using a reverse phase column (75 µm ID x 400 mm New Objective, in-house packed with ReproSil Gold 120 C18, 1.9 µm, Dr. Maisch GmbH) across a gradient from 5 to 40% in 38 min and 40 to 50% in 5 min (buffer A: 0.1% (v/v) FA; buffer B: 0.1% (v/v) FA, 95% (v/v) ACN). A targeted scheduled method was set up with one MS1 scan (R=70’000 at 400 m/z, normalized AGC =3e6 and max. IT 100 ms) and a cycle time of 3 s to monitor 57 peptides including iRT peptides (R=35’000 at 400 m/z, normalized AGC =1e5 and max. IT 110 ms) using an isolation window of 2.7 m/z and normalized collision energy 27%.

##### Statistical data analysis and visualization

For targeted analysis, peptides were analyzed manually using Skyline daily (Pino et al., 2020) and correct identification with at least 6 fragment ions per peptide was assigned based on the coelution profile of precursor and product ions. Peptide abundance was analyzed by summing the integrated areas of 6 fragment ions per peptide. Null values were imputed using random sampling from a normal distribution generated from 1% least intense values. Significance of change in intensity (ratio relative to total samples) was estimated using ROTS statistical test (ROTS R package, (Suomi et al., 2017)). Peptides were classified as non-amyloid, constitutive, pH-sensing reversible or irreversible amyloids based on arbitrary amyloid threshold of 0.715 relative intensity and significant differences (FDR ≤0.05 for ThT binding assay or ≤0.1 for pelleting assay) in distinct pH conditions. Analogically the ThT assay results were analyzed and classified using arbitrary threshold of 500 a.u. fluorescence intensity. Data was further visualized and analyzed with R 4.5. The entire dataset, including raw data, generated tables and scripts for the data analysis, is available in the PRIDE repository (ProteomeXchange with identifier PXD076452).

##### Comparing the TAPAAS output with bioinformatic predictions

Accuracy was defined as proportion of all correct classifications to total classifications. True positives rate (TPR) was defined as proportion of all correctly classified positives to all actual positives. False positives rate (FPR) was defined as proportion of all negatives that were incorrectly predicted as positives to all actual negatives, and for better distinction, split between FPR for non-amyloids and constitutive amyloids. Precision was defined as a proportion of all correctly classified positives to all predicted as positives.

#### Negative-staining transmission electron microscopy (TEM)

Peptide fibrillation was initiated using a stock solution of ARA1, Asn1_core_^WT^, Asn1_core_^non^, Asn1_core_^agg^ from peptides&elephants GmbH, were dissolved in dimethyl sulfoxide (DMSO; Sigma-Aldrich, D5879) at a concentration of 10 mg/ml and stored at −20°C until use. For formation: peptides were incubated at pH 5.0 or pH 7.5 in CPS buffer at a concentration of 0.5 mg/ml at 30°C for 24 h in low-binding Eppendorf Protein LoBind Tubes (Eppendorf, 0030 108 442). For reversibility: peptides were pre-incubated in CPS buffer at pH 5.0 at a concentration of 0.5 mg/ml at 30°C for 24 h in low-binding Eppendorf Protein LoBind tubes. Following pre-incubation, samples were divided into two tubes and centrifuged at 21’000 × g for 10 min at 25°C. The supernatant from both tubes were removed, and pellets were resuspended in CPS buffer at either pH 5.0 or pH 7.5. Samples after incubation were incubated for 1 min on carbon film 300-mesh copper grids (CF300-CU, Electron Microscopy Sciences). Samples were manually blotted using Whatman filter paper and washed twice with the same pH CPS buffer. The grid was negatively stained with 15 µl of 2% uranyl acetate and air-dried for 5 min. Morgagni 268 transmission electron microscope (Thermo Fisher Scientific) operating at 100 kV, 11,000x magnification, pixel size 1.64534 nm was used for collecting TEM images.

#### NMR data acquisition and analysis

For NMR measurements (n=1 per condition), high purity peptides with -CONH2 C-terminus were obtained from peptides&elephants GmbH, and diluted in DMSO-d_6_ at 10 mg/ml. For the pH titration, peptide samples were prepared at 0.5 mg/ml in CPS buffer at desired pH, supplemented with final 10% of a 1.1 mM DSS solution in D_2_O and measured immediately after preparation, sequentially from low pH to high pH. Spectra were recorded at RT (calibrated with methanol-d_4_, (Karschin et al., 2022)) in 3 mm diameter NMR tubes (Bruker). 1D ^1^H spectra using excitation sculpting for water suppression (Hwang and Shaka, 1995) were recorded on a 500 MHz AVNEO spectrometer equipped with a QCI-P cryo-probe (Bruker). The ^1^H spectral width was 15.6 ppm and 11k points were recorded in 256 scans, resulting in a measurement time of 7:30 min per sample. Spectra were processed and analyzed with the program Topspin 4.5.0 (Bruker). Spectra were calibrated with DSS and histidine chemical shifts were manually extracted. For measuring the signal intensity of the free peptide, the region from 0.6–1.0 ppm containing methyl signals was integrated and normalized to the intensity of the signal at the highest measured pH value.

#### Circular dichroism (CD) measurements

Peptides ARA1 or Asn1_core_^WT^ from peptides&elephants GmbH were dissolved in 100% DMSO at a concentration of 10 mg/ml and stored at −20°C until further use. For formation samples, 24 h before measurement peptides were diluted to 2 mg/ml in sterile-filtered CPS buffer and incubated at 30°C. For recovery samples, the formation process was started 48 h prior to measurement, the sample was split at 24 h prior to measurement, centrifuged (10 min, 14’000 × g) and the pellet resuspended into CPS pH 5.0 and pH 7.5 respectively, before incubation at 30°C was continued. Shortly before CD measurement, peptides were centrifuged (10 min, 14’000 × g) and washed in 1xPBS. To record CD spectra, quartz cuvettes (HellmaAnalytics) with 1mm path length were used in a J-815 CD Spectrometer (Jasco).

#### SAXS

High quality peptides (purity 95%) were ordered from peptides&elephants GmbH, dissolved in DMSO to a concentration of 10 mg/ml and stored at −20°C until further use. Peptides were diluted to a final concentration of 0.5 mg/ml in sterile-filtered CPS buffer at the stated pH and incubated for ∼24 h at 30°C before measurements in the Eppendorf Protein LoBind tubes (Eppendorf, 0030 108 442). Synchrotron EMBL P12 beam line at PETRA III (DESY, Hamburg, Germany) using a Pilatus 6M detector at a sample-detector distance of 3 m and at a wavelength of λ = 0.124 nm (I(q) vs q, where q = 4πsinθ/λ, and 2θ is the scattering angle) was applied for the measurements of the ARA1 and Asn1_core_^WT^ in CPS buffer as described above. 40 successive 0.1 s (4 s total exposure) frames were collected. We focus the analysis on the q-range 0.01 - 0.6 Å^−1^, since the data at low q’s might be influenced by the parasitic scattering from the beamstop or larger aggregates present in solution. The form factor fittings were carried out with SASView software using a scattering length density (SLD) of 14×10^−6^/ Å^2^ for the peptides and of 9.4×10^−6^/ Å^2^ for the solvent (Lattanzi *et al*., 2021).

### Analysis of full-length proteins *in vitro*

#### Protein purification

All bacterial strains used are listed in the key resource table. *E. coli* Rosetta pLys cells were transformed with plasmids expressing Asn1_FL_^WT^ (kindly provided by Kollman laboratory) wild-type or mutants Asn1_FL_^non^ (mutation introduced via mutagenesis with primers into Asn1_FL_^WT^). *E. coli* ArcticExpress (DE3) (Agilent, 230192) cells were transformed with plasmids expressing Asn1_FL_^agg^. Cells were grown at 37°C (ex. 30°C for Asn1_FL_^agg^) in TB media (23.6 g/l yeast extract, 11.8 g/l tryptone, 9.4 g/l K_2_HPO_4_, 2.2 g/l KH_2_PO_4_, 0.4% glycerol) with kanamycin, and protein expression was induced at OD_600_=1.4 with 0.1 mM IPTG (ex. OD_600_=1 for Asn1_FL_^agg^). After growth at 16°C for 24 h (ex. 10°C for 30 h for Asn1_FL_^irrev^), cells were harvested by centrifugation and the pellet resuspended in lysis buffer (50 mM NaPO_4_, 500 mM NaCl, 20 mM imidazole, pH 8.0) supplemented with cOmplete™, Mini, EDTA-free Protease Inhibitor Cocktail (Roche, 11836170001), DNase I recombinant (Roche, 04536282001), 0.5 mg/ml Lysozyme Biotin-Caproyl (Sigma-Aldrich, L0289). Cells were lysed for 1 h in 4°C. Cell disruption was performed using an EmulsiFlex homogenizer by passing the suspension through the instrument at high pressure until complete lysis was achieved. The lysate was then clarified by centrifugation in the at 20’000 × g for 60 min at 4°C. Protein purification was performed using an ÄKTA chromatography system. The supernatant was loaded at 4°C onto a HisTrap™ HP His tag 5 ml (Cytiva, 17524802) column following the manufacturer’s instructions, proteins eluted with elution buffer (50 mM NaPO_4_, 500 mM NaCl, 500 mM imidazole pH 8.0). Eluted fractions were collected and analyzed by SDS-PAGE. Selected fractions were pooled and loaded onto a gel filtration column HiLoad^®^ 16/600 Superdex^®^ 200 pg (Cytiva 28-9893-35) equilibrated with gel filtration buffer (20 mM Tris, 150 mM NaCl, 5 mM DTT, pH 7.5). Collected fractions were analyzed by SDS-PAGE and Coomassie blue staining. Pure proteins were aliquoted and stored at −80°C.

#### Microfluidics (PhaseScan)

##### Protein Labelling

Purified proteins were thawed on ice and clarified by centrifugation at 21’000 × g for 10 min at 4°C. The soluble fraction was subsequently labelled with Alexa Fluor 647 NHS ester dye (Thermo Fisher Scientific). Briefly, soluble protein was incubated with the dye at a molar ratio of 1:2 for 25 min at room temperature, without agitation, and protected from light. Following incubation, excess free dye was removed using Zeba spin columns (Thermo Fisher Scientific) according to the manufacturer’s instructions. Columns were pre-equilibrated by two successive washes with CPS buffer prior to sample application. The labelled protein was eluted by centrifugation. Protein concentration was determined by measuring absorbance at 280 nm using a spectrophotometer, with correction for dye contribution at this wavelength. Finally, labelled protein was mixed with unlabelled protein to achieve a final labelling fraction of 1% and stored protected from light until further use.

##### Fabrication of microfluidic devices

The microfluidic devices were fabricated as described previously (Arter *et al*., 2022; Agarwal *et al*., 2025). Briefly, devices were designed in AutoCAD and produced using standard soft-photolithography. SU-8 3050 (A-Gas Electronic Materials Limited) was spin-coated onto polished silicon wafers (MicroChemicals GmbH) to a height of ∼50 µm, soft-baked, UV-exposed through a transparency mask, post-baked, and developed in propylene glycol monomethyl ether acetate (PGMEA, Sigma-Aldrich) to generate the master mould. Poly(dimethylsiloxane) (PDMS Sylgard 184 kit; Dow Corning, 10:1 base:crosslinker) was cast on the master, cured at 60°C for 2 h, cut to shape, and inlet/outlet ports were punched. After sonication in isopropanol, PDMS devices were bonded to glass slides (Epredia) via oxygen plasma (60% power for 30 s, Diener Femto Electronics). Channel surfaces were made hydrophobic by treatment with 1% (v/v) trichloro(1H,1H,2H,2H-perfluorooctyl)silane (Sigma) in (HFE-7500 3M™ Novec™ Engineered fluid), followed by drying at 95°C.

##### Generation of phase diagrams

PhaseScan, a semi-automated combinatorial droplet microfluidic platform, was used to construct multidimensional phase diagrams, as described previously (Arter *et al*., 2022; Agarwal *et al*., 2025). Briefly, each PhaseScan experiment consisted of aqueous inputs: protein, pH 4 buffer, and pH 8 buffer. The acidic and basic buffers were prepared using 200 mM citric acid buffer, 200 mM sodium phosphate buffer containing 150 mM NaCl at pH 4 and pH 8, respectively. Pressure-driven flow control was implemented using Flow EZ™ controllers (Fluigent) to deliver the aqueous streams and fluorinated oil to the droplet generator. Each aqueous solution carried a unique fluorescent barcode: protein was labelled with Alexa Fluor 647 NHS ester (1% labelling), the pH 4 buffer contained free Alexa Fluor 488, and the pH 8 buffer contained free Alexa Fluor 546 (Thermo Fisher Scientific). Droplet pH values were inferred from a calibration curve generated by mixing the acidic and basic buffers at defined ratios (Ausserwöger *et al*., 2026). Automated flow profiles varied the relative aqueous inputs to generate a continuous concentration landscape. HFE-7500 oil containing 1.2% (w/v) fluorosurfactant (RAN Biotechnologies) was supplied at a constant rate. Droplets were imaged under continuous flow using an epifluorescence microscope equipped with a 10× objective (Cairn Research). Image analysis was performed with custom Python scripts. Droplets were segmented, quality-filtered, and their barcode intensities quantified and normalized to droplet volume before conversion to absolute concentrations using calibration standards. Droplets were classified as homogeneous or with aggregates based on the presence of condensates, and the resulting data were visualized as scatter plots representing mean values across conditions.

#### Pelleting assay

Purified proteins Asn1_FL_^WT^, Asn1_FL_^non^ or Asn1_FL_^agg^ were thawed on ice, cleared by centrifugation (4°C, 10 min, 21’000 × g), and diluted to a final protein concentration of 5 µM in CPS buffer with additional 0.2 mM TCEP at either pH 5.5 or pH 7.5. Samples were incubated at 25°C for 24 h in a CLARIOstar Plus plate reader (BMG LABTECH) shaking at 200 rpm. A 15 µL sample was collected for the total analysis, before aggregates were pelleted by centrifugation (4°C, 10 min, 21’000 × g), and separated from the supernatant containing soluble protein. Supernatant fraction of 15 µL were collected. The pellet was resuspended in cracking buffer (8 M urea, 50 g/l SDS, 40 mM Tris-HCl [pH 6.8], 0.1 mM EDTA, 0.4 mg/ml Bromophenol Blue, 2% b-Mercaptoethanol). All samples were mixed with sample buffer and incubated at 95°C for 5 min. 15 µl of each sample was separated on 4-12% Bis-Tris gels (Invitrogen, NW04127) running in MOPS SDS buffer. Gels were stained with Coomassie blue, bands analyzed with FIJI and quantified as gray measured area normalized to the empty gel background.

#### Cryo-EM analysis

##### Sample preparation

Purified protein Asn1 was adjusted to a final concentration of 20 µM in CPS buffer adjusted with HCl to pH 5.5. The sample was incubated in a CLARIOstar Plus plate reader (BMG LABTECH) at 25°C with continuous shaking at 200 rpm for 24 h. Following incubation, the protein solution was supplemented with 0.05% (w/v) optoglucoside. After incubation, 3.5 µl of the sample was applied to glow-discharged (25 mA for 30 s using a PELCO easiGlow) Quantifoil R1.2/1.3 Cu 300-mesh grids that had been manually pre-coated with 2.3 nm non-continuous carbon. The sample was plunge-frozen using a Vitrobot IV (Thermo Fisher Scientific) at 8°C and 95% humidity, with a blot time of 4.5 s, a wait time of 60 s, and a blot force of 0 (Tivol et al. 2008).

##### Cryo-EM data collection

Cryo-EM data of Asn1 filaments from purified protein were collected at the ETH Zurich ScopeM facility on a Titan Krios G4 equipped with a Gatan GIF BioContinuum energy filter, operated at a slit width of 20 eV and K3 camera (Gatan), operated at counted mode. The data were collected at a nominal magnification of 81’000× (1.06 Å/pix) with EPU software (Thermo Fisher Scientific, version 3.10.0) with a total exposure time of 2.0 s and an accumulated dose of 48 e⁻/Å². The defocus range was set from −1.2 to −2.8 µm with an incremental step of 0.2. A total of 11’312 movie stacks, each consisting of 40 frames, were collected (Table S11).

##### Cryo-EM SPA processing

The image analysis was done in CryoSPARC 5.0.2 (Punjani *et al*., 2017) using a standard pipeline. The processing scheme of Asn1 is shown in Fig. S5 and Table S11. The gain reference was created *a posteriori* (Afanasyev *et al*., 2015) from 40’000 frames and used in patch motion-correction. The statistics of the results, together with the CTF estimation results, were used in data curation and selection of the best micrographs. A reference-free filament tracer was used for picking an initial set of particles at a separation distance of 100 Å. The particles were extracted at a resulting pixel size of 11.65 Å/pix and subjected to 2D-classification. The 2D class-averages showed strong compositional heterogeneity of the filaments, with clear difference in their thickness. Particles from various 2D class-averages were subjected to a multi-class *ab initio* reconstruction, yielding three distinct 3D-classes, corresponding to various filament populations composed of 7, 5, and 4 protofilaments (3D-class 1-3, respectively). One “bad” 3D-class was also reconstructed from the same particle stack. Particles corresponding to the 3D-class 3 were selected and subjected to 2D-classification. The resulting 2D class-averages were used as templates for template-based particle re-picking, yielding 717’737 particles. These particles were subjected to 2D-classification, and only classes displaying well-defined 2D class-averages (221’535 particles) were retained for heterogeneous refinement. The particles were extracted at a pixel size of 2.12 Å/pix and subjected to heterogenous refinement to remove particles corresponding to 3D-classes, composed of more than 4 protofilaments. The resulting particle stack (134’334 particles) corresponding to the 3D-class 3 was further refined with non-uniform refinement. Next, symmetry expansion (C4) was applied followed by local refinement focused on one strand of the filament (537’336 particles). Then, each asymmetric unit of the reconstruction was recentered and aligned relative to each other. The resulting particle stack of 1’074’672 particles was re-extracted at 1.06 Å/pix and locally refined, resulting in an estimated resolution of approximately 4.8 Å.

Cryo-EM density maps for mask generation were created in ChimeraX (Goddard *et al*., 2018) using the “Cube’n Tube” plugin (Cairoli T. *Cube’n Tube* Version 1.1.0, 2026, DOI 10.5281/zenodo.18754998).

##### Model building and refinement

An initial atomic model of the Asn1 subunit was predicted using AlphaFold3 (Abramson *et al*., 2024). The protein was rigid-body fit into the cryo-EM density map. The model refinement was performed in PHENIX (Liebschner *et al*., 2019) using phenix.real_space_refinement (Afonine, Poon, *et al*., 2018) as iterated in Table S11. The refined model fit was manually inspected in COOT (Casañal, Lohkamp and Emsley, 2020). The resulting model was validated using MolProbity (Williams *et al*., 2018) and the available tools in PHENIX (Afonine, Klaholz, *et al*., 2018).

Figures were prepared using ChimeraX and Adobe Illustrator. The corresponding cryo-EM map was deposited in the Protein Data Bank under accession code (PDB ID: 31ZZ).

##### *In vitro* activity measurements

Purified proteins Asn1_FL_^WT^, Asn1_FL_^non^ or Asn1_FL_^agg^ were thawed on ice, cleared by centrifugation (4°C, 10 min, 21’000 × g), and diluted to a final protein concentration of 5 µM at pH 5.5 or pH 7.5 in CPS buffer with additional 0.2 mM TCEP (total sample volume is 100 µL). Samples were incubated in CLARIOstar Plus plate reader (BMG LABTECH) 200 rpm, 25°C, 24 h. The enzymatic activity of proteins Asn1_FL_^WT^ vs Asn1_FL_^non^ or Asn1_FL_^agg^ was assessed by measuring the production of AMP with a commercially available luminescence assay kit AMP-Glo™ Assay (Promega, V5012), as described previously (Chang *et al*., 2023). Enzyme reactions were prepared in duplicate by pipetting 12.5 µl of protein (100 nm of each) diluted in CPS buffer at pH 5.5 or pH 7.5 into PCR tubes. The reaction was initiated with the addition of 12.5 µl substrate solution (CPS pH 5.5 or CPS pH 7.5, 10 mM MgCl_2_, 0.75 mM DTT, 0.05 mg/ml BSA, 10 mM aspartate, 10 mM glutamine, and 1 mM ATP), well mixed, centrifuged and incubated 0, 5, 15, 30 min at RT. After the indicated incubation time, the reaction was quenched by adding 25 µl of AMP-Glo^©^ I to each reaction tube, well mixed, centrifuged and incubated 1 h at RT. Next, 50 µl of AMP detection solution (AMP-Glo^©^ Reagent II + Kinase Glo^®^ One Solution) was added, well mixed, centrifuged and incubated 1 h at RT. Samples were transferred into a 96 well plate with solid bottom (Milipore, MSSWNFX40) and read out in a CLARIOstar Plus (BMG LABTECH, 0.2 s, scanning diameter 4 mm, emission 521-20, gain 3’600, focal height adjusted, mixed 30 s 300 rpm) plate reader. Luminescence of each well was determined as relative luminescent units. Quantities of produced AMP were ascertained by referencing an AMP standard curve covering 0 µM −10 µM buffer for each pH independently (CPS pH 5.5 or CPS pH 7.5, 10 mM MgCl_2_, 0.75 mM DTT, 0.05 mg/ml BSA, and 10 mM ATP).

### *In vivo* and *ex vivo* yeast methods

#### Yeast strains

Yeast strains are listed in Table S12. Yeast strains (Asn1-mNG, Asn2-mNG, Bdh1-mNG, Gad1-mNG, Dcs1-mNG) were constructed with introduction mNG tag via homological recombination into WT yeast BY4741(Lys2D0Met15DLeu2D0 /a) with pFA6a-mNeonGreen-HIS3MX6 (Addgene, Plasmid #129100). Asn1^WT^-mNG was introduced into pSIV-URA and mutations introduced into pSIV-URA-Asn1^WT^ by site directed mutagenesis via primers. Plasmid pSIV-URA-Asn1^WT^ or pSIV-URA-Asn1^mut^ was linearized and inserted into URA locus via homological recombination. Other strains were constructed via mating and dissection. Deletion strains were constructed with Kan using pFA6a-mNeonGreen-KanMX6 (Addgene, Plasmid #129099) cassette.

#### Yeast cell growth and fluorescence microscopy

Cells were grown in synthetic SD media (2% glucose, 0.5% NH4-sulfate, 0.17% yeast nitrogen base, and specific amino acids i.e. for minimal media: HLMK and uracil (named -ASN) and if stated asparagine (named +ASN)) at 30°C. To analyze exponentially growing cells, overnight cultures were diluted to OD_600_=0.2 and incubated at 30°C until reaching logarithmic growth OD_600_=0.4-0.6. For stationary phase cultures, the diluted cells were incubated at 30°C for 3 days before analysis. For observation of growth, cells were either spotted in 10x serial dilutions on SD agar (named -ASN or +ASN) plates and the plates were imaged after 3 days at 30°C or cells were loaded at OD_600_ 0.1 into 24-well plates (Thermo Scientific Nunclon Delta Surface Cat.142475) and optical density at 600 nm was analysed using a Spectrostar plate reader. Fluorescence microscopy was performed using a Nikon Eclipse Ti-E microscope with NIS-Elements Advanced Research V 5.02 software (for mNG strains: GFPdual channel, for Ded1-mCherry: mCherrydual channel, 200 ms exposure, 49% and 73% laser power, respectively, 5 µm z-stack a 27 slices). Microscopy experiments were performed in triplicates, unless stated otherwise. Maximal intensity projection images were generated using imageJ software. Cells with foci were counted manually. For time-lapse experiments, yeast cells were loaded in commercial microfluidic chips (CellASIC ONIX2, Merck Millipore) and images were recorded every 10 min.

#### Microscopy screen

Yeast strains from the GFP collection (see Suppl. Table S12 for all strains, (Huh *et al*., 2003)) were inoculated overnight in SD complete media in 96-well sterile plates (Thermo Scientific, 249946) and covered with gas permeable membranes (Sigma-Aldrich, Z380059), 30°C, 230 rpm. In the morning diluted to OD_600_=0.1-0.2, grown at 30°C, 230 rpm were imaged during exponential growth, at 2 days stationary phase or 1h after re-addition of fresh media, using Glass Clear Bottom Black 96 Well Plate (Matrical MGB096-1-2-LG-L). Fluorescence microscopy was performed using a Nikon Eclipse Ti-E microscope with NIS-Elements Advanced Research V 5.02 software (strains: GFPdual channel, 100ms exposure 2 µm z-stack, 9 Z-positions). Cells with foci were counted manually in FIJI, Z projection, MaxIntensity. Proteins were classified based on the percentage of cells containing GFP foci in log phase, 2-day stationary phase, and recovery. Constitutive proteins showed foci in >10% of cells in stationary phase and recovery, and >5% in log phase. Reversible proteins showed foci in >10% of cells in stationary phase, but <5% in log phase and <10% after recovery. Non-foci proteins showed low foci formation in all conditions: <10% in stationary phase, <5% in log phase, and <10% after recovery. Partially reversible proteins were the remaining cases where the percentage of cells with foci decreased after recovery compared with stationary phase.

#### SDD-AGE-MS

##### Sample preparation

Yeast cells were grown in SD complete media containing all amino acids overnight, diluted to OD_600_=0.2 and collected as exponentially growing (OD_600_=0.6-0.7) or 6 day stationary phase growth 30°C, totally OD_600_=25 cell pellet was collected for every sample. For protein extraction, pellets were resuspended in approximately 300 µL of ice-cold lysis buffer containing 50 mM Tris-HCl pH 7.5, 150 mM NaCl, 1% (v/v) Triton X-100, 2.5 mM EDTA, N-ethylmaleimide (NEM), phenylmethylsulfonyl fluoride (PMSF), and 1× protease inhibitor cocktail. Ice-cold glass beads were added to the suspension, and cells were lysed by mechanical disruption at 6 m/s for three cycles of 30 s each, with 5 min cooling intervals between cycles. Lysates were clarified by centrifugation, and the resulting supernatants were used for semi-denaturing detergent agarose gel electrophoresis (SDD-AGE). Protein concentrations were determined using a bicinchoninic acid (BCA) assay according to the manufacturer’s instructions (Thermoscientific, Pierce BCA Protein Assay, 23227). Supernatants were adjusted to equal protein concentrations prior to electrophoresis. Normalized lysates were mixed with 4× sample buffer (40 mM Tris-acetate, 2 mM EDTA, 20% glycerol, 4% SDS, and bromophenol blue). Samples were incubated for 10 min at room temperature before loading. For each lane, approximately 20 µl of sample (depending on the concentration) was prepared together with 5 µl of GeneRuler marker (Thermo Scientific, GeneRuler 1 kb DNA Ladder, SM0311), which was added immediately before loading. SDD-AGE gels were cast at 1.0 mm thickness using 1% agarose in 1× TAE (40 mM Tris-acetate and 1 mM EDTA) supplemented with 0.1% SDS. Electrophoresis was carried out at 60 V for 1 h 15 min at 4°C. After electrophoresis, gels were incubated in 15 ml water containing 5 µl of intercalating dye Gel Red (Biotium, 41003) and the DNA GeneRuler marker was visualized under a UV transilluminator. Five fractions were cut at the following marker lanes: 10’000 bp, 3’000 bp, 1’000 bp and 250 bp, excised and dried using a speedvac device. Once the samples were dried, these were resuspended for 30 min at 37°C in 100 µl of a solution composed of ammonium bicarbonate (50 mM) and TCEP (5 mM). 10 mM iodoacetamide was added to the solution (30 min, 37°C in the dark). The supernatant was removed and the agarose bands were dried in the speedvac device and resuspended in 100 µl of 50 mM ammonium bicarbonate and proteolyzed with 1 µg of trypsin (overnight, 37°C). The following day the proteolysis was quenched with 5% formic acid, subjected to C18 cleanup (MiniSpin, The Nest group) and resuspended in 20 µl of MS buffer (2% acetonitrile, 0.1% formic acid) and 8 µl were subjected to LC-MS analysis.

##### MS acquisition

LC–MS/MS analyses were conducted using an Orbitrap Q Exactive Plus mass spectrometer (Thermo Fisher Scientific) interfaced with an EASY-nLC 1000 liquid chromatography system (Thermo Fisher Scientific). Peptide separation was achieved on a reversed-phase analytical column (75 µm inner diameter × 400 mm length; New Objective), in-house packed with ReproSil Gold 120 C18 resin (1.9 μm particle size; Dr. Maisch GmbH). Peptides were eluted using a gradient of 5–25% buffer B over 55 min, followed by an increase from 25–40% buffer B over 5 min (buffer A: 0.1% (v/v) formic acid; buffer B: 0.1% (v/v) formic acid, 95% (v/v) acetonitrile). Data were acquired in data-dependent acquisition (DDA) mode, with one MS1 scan followed by up to 20 MS/MS scans of the most intense precursor ions. MS1 spectra were acquired at a resolution of 70’000 (at m/z 400), with an automatic gain control (AGC) target of 3 × 10⁶ and a maximum injection time of 64 ms. Peptide fragmentation was performed using higher-energy collisional dissociation (HCD) with a collision energy of 25% and an isolation window of 1.4 m/z. MS2 spectra were acquired at a resolution of 17,500 (at m/z 400), with an AGC target of 1 × 10⁵ and a maximum injection time of 55 ms. Dynamic exclusion was set to 30 s, and precursor ions with charge states below 2 or above 6 were excluded from selection.

##### Data analysis

Acquired spectra were searched using the MaxQuant software package version 2.0.3.0 embedded with the Andromeda search engine (PMID: 19029910) against the respective reference dataset (http://www.uniprot.org/, downloaded on 17.05.2021, 6’057 proteins) extended with reverse decoy sequences. The search parameters were set to include only full tryptic peptides, maximum two missed cleavage, carbamidomethyl as static peptide modification, oxidation (M) as variable modification (maximum 5 modifications per peptide) and “match between runs” option disabled. The MS and MS/MS mass tolerance was set to 10 ppm, and a false discovery rate at PSM and protein level of 0.01. Protein abundance was determined (MaxLFQ intensities) from the intensity of top two peptides. Proteins identified in fraction 5 in at least two replicates per condition were considered. Subsequently, the protein intensity of fraction 5 (highest molecular weight) was normalized with the sum of the intensity of all fractions. Null values were imputed using random sampling from a normal distribution generated from 5% less intense values. Protein intensities were compared between two experimental conditions (n=3). The statistical testing was performed using the ROTS algorithm (Suomi et al., 2017) with default 5’000 bootstrap resampling and K equal to the number of tested proteins. Overlap between proteins enriched at higher molecular weight (upper band) during cell starvation (log₂FC > 1, FDR ≤ 0.1) and various reference datasets curated from the indicated literature was evaluated using a hypergeometric test (R standard package, p value adjusted with Holm correction). GO terms enrichment analysis was performed as described above. The entire dataset, including raw data and generated tables, is available in the PRIDE repository (ProteomeXchange with identifier PXD077539).

#### pH shifts in stationary phase

Yeast cells expressing potential candidates fused to an mNeonGreen tag were grown to stationary phase for 72 h in SD complete media. The resulting stationary phase cultures were pelleted at RT at 21’300 rcf for 2 min and the pH of the supernatant was either kept non-pH-adjusted or adjusted to pH 7.5 with 5 M NaOH and subsequently filter sterilized. Cells were then loaded in commercial microfluidic chips (CellASIC ONIX2, Merck Millipore), released into the non-adjusted or adjusted media and images were recorded every 5 min as described above and cells with foci were counted manually.

#### Potassium sorbate treatment

Yeast cells were grown to exponential phase in SD complete media as described above. Cells were immobilized on 96-well plate using concanavalin A and media was exchanged for SD complete media adjusted to pH 4.5 and supplemented with 0 mM, 2 mM, 4 mM, 6 mM or 8 mM potassium sorbate to reduce intracellular pH as described in (Joyner *et al*., 2016). The plates were incubated for 30 min at 30°C at 120 rpm. Cells were imaged as described above and cells with foci were counted manually.

#### Western Blot

Stationary phase cells grown in minimal media for 3 days were harvested and adjusted to OD₆₀₀=1. Pellets were washed by centrifugation, residual medium removed, and protein samples were prepared by TCA extraction and acetone precipitation. 15 µl of lysate was separated on bolt 4-12% Bis-Tris gels (Invitrogen, NW04127) in MOPS SDS buffer followed by standard western blotting with primary antibodies anti-mNeonGreen 1:3’000 (mNeonGreen Polyclonal antibody Protein Tech group 29523-1-AP), and anti-Pgk1 1:3’000 (anti-Pgk1, mouse IgG1, monoclonal 22C5, Molec. Probes 459250) Blots were developed with Clarity Western ECL Substrate solutions (BioRed, 1705061) and scanned on a Fusion FX7 imaging system (Witec AG). Blots were quantified via FIJI as a gray mean value. mNG relative intensity levels were normalized to the Pgk1 ratio.

