## Supplementary information for "Systematic identification of pH-sensing amyloid core motifs reveals a widespread mechanism for reversible protein assembly upon stress"

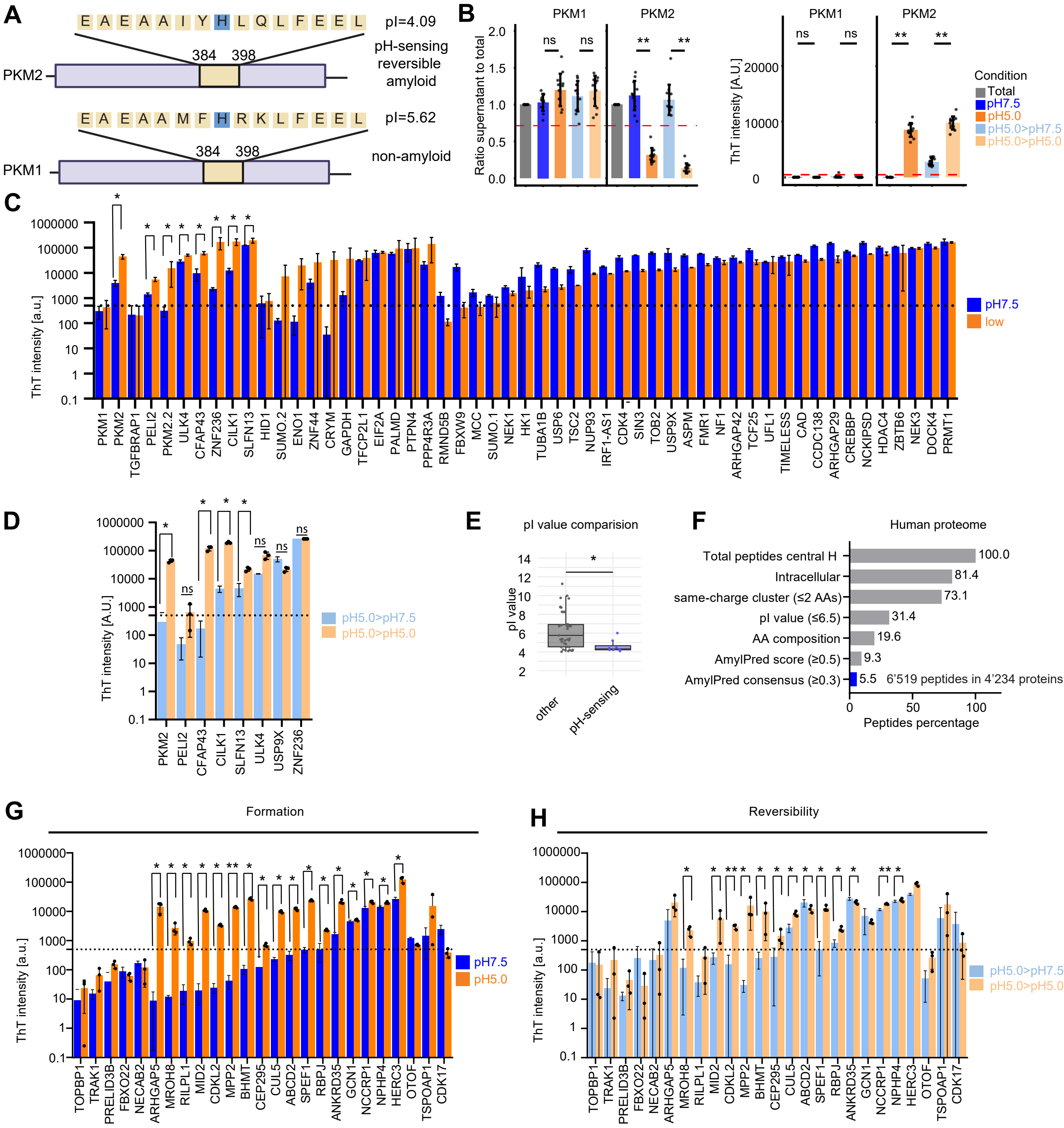
A
E A E A A I Y H L Q L F E E L
pI=4.09
pH-sensing reversible amyloid
PKM2
384 398
E A E A A M F H R K L F E E L
pI=5.62
non-amyloid
PKM1
384 398
B
PKM1
PKM2
Ratio supernatant to total
ns
**
ThT intensity [A.U.]
Condition
Total
pH7.5
pH5.0
pH5.0>pH7.5
pH5.0>pH5.0
C
ThT intensity [a.u.]
pH7.5
low
PKM1 PKM2 TGFBRAP1 PELI2 PKM2.2 ULK4 CFAP43 ZNF236 CILK1 SLFN13 HID1 SUMO.2 ENO1 ZNF44 CRYM GAPDH TFCP2L1 EIF2A PALMD PTPN4 PPP4R3A RMND5B FBXW9 MCC SUMO.1 NEK1 HK1 TUBA1B USP6 TSC2 NUP93 IRF1-AS1 CDK4 SIN3 TOB2 USP9X ASPM FMR1 NF1 ARHGAP42 TCF25 UFL1 TIMELESS CAD CCDC138 ARHGAP29 CREBBP NCKIPSD HDAC4 ZBTB6 NEK3 DOCK4 PRMT1
D
ThT intensity [A.U.]
pH5.0>pH7.5
pH5.0>pH5.0
PKM2 PELI2 CFAP43 CILK1 SLFN13 ULK4 USP9X ZNF236
E
pI value comparision
pI value
other
pH-sensing
F
Human proteome
Total peptides central H 100.0
Intracellular 81.4
same-charge cluster (≤2 AAs) 73.1
pI value (≤6.5) 31.4
AA composition 19.6
AmylPred score (≥0.5) 9.3
AmylPred consensus (≥0.3) 5.5
6'519 peptides in 4'234 proteins
Peptides percentage
G
Formation
ThT intensity [a.u.]
pH7.5
pH5.0
TOPBP1 TRAK1 PRELID3B FBXO22 NECAB2 ARHGAP5 MROH8 RILPL1 MID2 CDKL2 MPP2 BHMT CEP295 CUL5 ABCD2 SPEF1 RBPJ ANKRD35 GCN1 NCCRP1 NPHP4 HERC3 OTOF TSPOAP1 CDK17
H
Reversibility
ThT intensity [a.u.]
pH5.0>pH7.5
pH5.0>pH5.0
TOPBP1 TRAK1 PRELID3B FBXO22 NECAB2 ARHGAP5 MROH8 RILPL1 MID2 CDKL2 MPP2 BHMT CEP295 CUL5 ABCD2 SPEF1 RBPJ ANKRD35 GCN1 NCCRP1 NPHP4 HERC3 OTOF TSPOAP1 CDK17

**Supplementary Figure S1: Establishing criteria to predict and experimentally validate pH-sensing, reversible amyloid peptides**

A. Cartoon representation of the human pyruvate kinases PKM1 and PKM2, highlighting the amyloid core region with the predicted pI value. Note that the amyloid core motif of PKM2, but not PKM1, assembles pH-sensing, reversible amyloids (sequences adapted from (Cereghetti *et al.*, 2024)).

B. TAPAAS sub-assay results of the 15-mer control peptides of non-amyloid PKM1 and pH-sensing, reversible amyloid PKM2 incubated at pH 7.5 or pH 5.0 for 24 h at 30°C. For reversibility, peptides were pre-incubated at pH 5.0, and then shifted to pH 7.5 or maintained at pH 5.0 for 24 h at 30°C. An aliquot before incubation was used as input control (Total). To quantitatively probe aggregation, relative peptide abundance in the supernatant was measured by mass spectrometry after pelleting (left) or ThT staining (right, in arbitrary units (a.u.)). Each point shows the mean of experimental triplicate values ($n \geq 11$) ± standard deviation (SD). The dashed lines indicate the chosen amyloid thresholds of 500 a.u. (ThT assay) or the 0.715 ratio of soluble peptide in the supernatant compared to the input (total) in the pelleting assays; pairwise Wilcoxon rank-sum test with Holm correction; ** $p < 0.01$; ns, not significant.

C. ThT binding of 51 initial peptides (concentration 2 mg/ml) with PKM1 and PKM2 peptide controls probing amyloid formation at pH 7.5 or low pH (in TRIS buffer, pH 5.8) for 24 h at 30°C; n=3, mean ± SD, a.u.: arbitrary units. The dotted line indicates arbitrary thresholds for amyloids at 500 a.u.; ROTS, * FDR <0.05 and $p < 0.05$; only significance for pH-sensing peptides is shown.

D. ThT binding probing reversibility of pH-sensing candidates from Fig. S1C at the concentration of 2 mg/ml. The amyloids were pre-formed at pH 5.0 for 24 h at 30°C, before the pH was switched to 7.5 or kept at 5.0 for control for an additional 24 h at 30°C; n=3, mean ± SD, a.u.: arbitrary units. The dotted line indicates the arbitrary threshold for amyloids at 500 a.u.; ROTS, * $p < 0.05$; ns, not significant.

E. Comparison of theoretical pI values for pH-sensing peptides compared to other tested peptides from panel C. A Wilcoxon rank-sum test controls statistical significance; * $p < 0.05$.

F. Sequential filtering criteria for pH-sensing amyloid motifs were applied to the human proteome and the percentage (%) of predicated peptides remaining after each step is shown. 6’519 peptides, representing 5.5 % of all 15-mer peptides with a central histidine and no prolines, from 4’234 proteins scored by these criteria as pH-sensing.

G. and H: ThT binding of 25 pH-sensing peptides (at 0.5 mg/ml) predicted from 1’777 candidates by the improved bioinformatic workflow (Fig. 1F) after incubation performed as in panels (C, D) at the indicated pH conditions probing formation (G) and reversibility (H); n=3, mean ± SD, ROTS, a.u.: arbitrary units, * FDR<0.05 and p <0.05, ** FDR <0.05 and p <0.01; statistical significance is only shown for pH-sensing peptides.

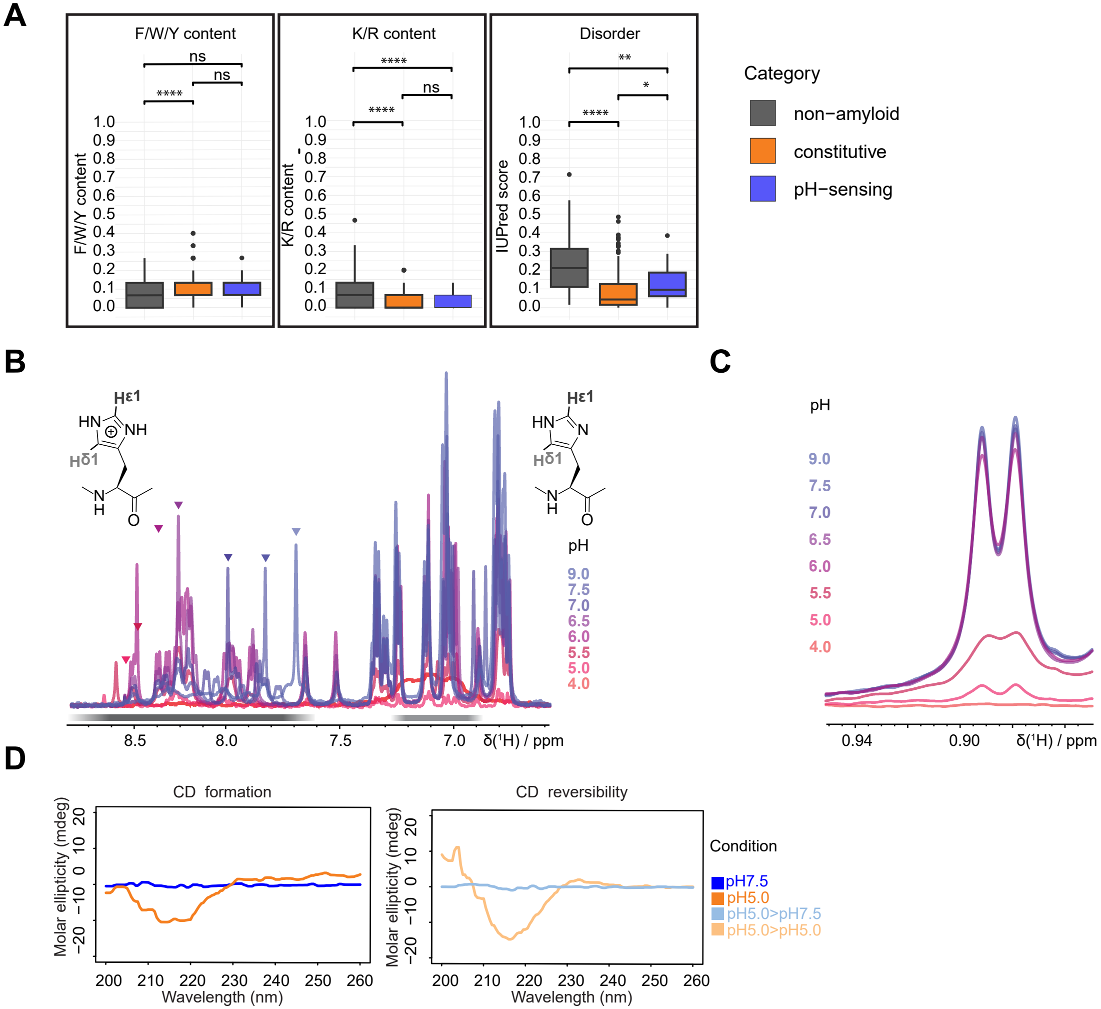
A
F/W/Y content
K/R content
Disorder
F/W/Y content
K/R content
IUPred score
ns
****
**
*
Category
non−amyloid
constitutive
pH−sensing
B
Hε1
Hδ1
HN
NH
N
H
O
pH
9.0
7.5
7.0
6.5
6.0
5.5
5.0
4.0
8.5
8.0
7.5
7.0
δ(¹H) / ppm
C
pH
0.94
0.90
δ(¹H) / ppm
D
CD formation
CD reversibility
Molar ellipticity (mdeg)
Wavelength (nm)
Condition
pH7.5
pH5.0
pH5.0>pH7.5
pH5.0>pH5.0

**Supplementary Figure S2: Biophysical properties of identified amyloid cores lead to design of artificial pH-sensing reversible amyloid peptide**

A. Comparison of selected properties including aromatic (F/W/Y) and positively charged (K/R) amino acid content, and disorder of experimentally validated pH-sensing (n=28), constitutive (n=156) and non-amyloid (n=125) peptides. The box plots show median, Q1-Q3 ± max 1.5 IQR and outliers, Kruskal-Wallis and Dunn post-hoc-test with Holm correction; * $p < 0.05$, ** $p < 0.01$, *** $p < 0.001$, **** $p < 0.0001$; ns, not significant.
B. Overlay of 1D $^{1}$H NMR spectra recorded across a pH gradient from 4 to 9, as indicated by the color code (right). The $^{1}$H resonance corresponding to $H^{\varepsilon 1}$ of the central histidine within the ARA1 peptide is indicated with triangles. The progressive chemical shift changes reflect protonation-dependent alterations of the histidine, which was used for the plot in Fig. 2E.
C. Excerpt of 1D $^{1}$H NMR spectra of representative methyl resonances of ARA1 recorded across a pH gradient from 4 to 9, as indicated by the color code (left). Disappearance of the signals reveals pH-dependent aggregation of the peptide.
D. Representative circular dichroism (CD) spectra of the ARA1 peptide recorded after 24 h incubation at 30°C at pH 5.0 or pH 7.5 (formation, left). To probe reversibility, ARA1 fibrils formed at pH 5.0 were analyzed after the pH was switched to 7.5 or kept at 5.0 for control for an additional 24 h at 30°C (right). Molar ellipticity (mdeg) as a function of wavelength is plotted (n=2).

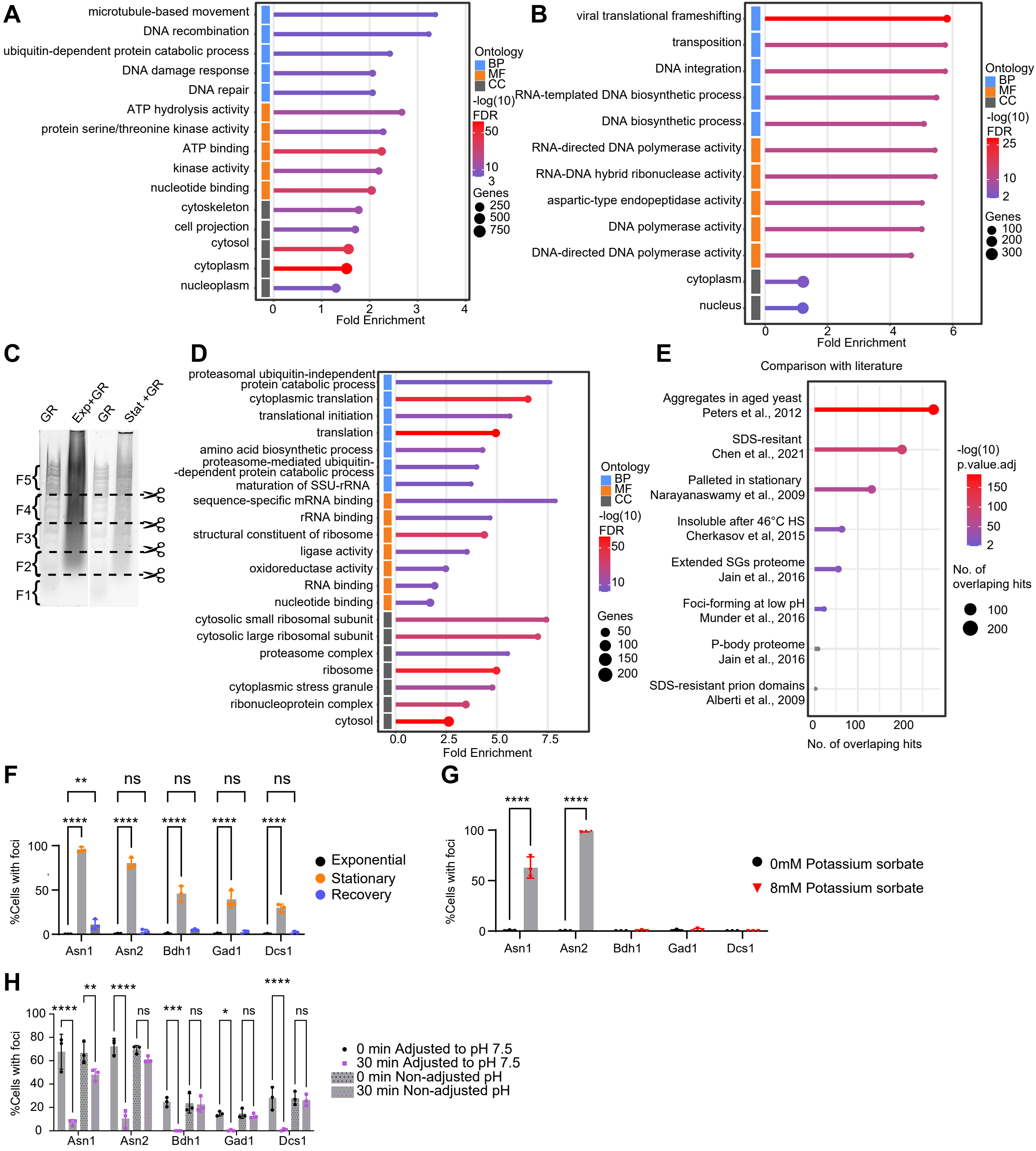

A
microtubule-based movement
DNA recombination
ubiquitin-dependent protein catabolic process
DNA damage response
DNA repair
ATP hydrolysis activity
protein serine/threonine kinase activity
ATP binding
kinase activity
nucleotide binding
cytoskeleton
cell projection
cytosol
cytoplasm
nucleoplasm
Fold Enrichment
Ontology
BP
MF
CC
-log(10) FDR
Genes
B
viral translational frameshifting
transposition
DNA integration
RNA-templated DNA biosynthetic process
DNA biosynthetic process
RNA-directed DNA polymerase activity
RNA-DNA hybrid ribonuclease activity
aspartic-type endopeptidase activity
DNA polymerase activity
DNA-directed DNA polymerase activity
cytoplasm
nucleus
Fold Enrichment
C
GR
Exp+GR
GR
Stat +GR
F5
F4
F3
F2
F1
D
proteasomal ubiquitin-independent protein catabolic process
cytoplasmic translation
translational initiation
translation
amino acid biosynthetic process
proteasome-mediated ubiquitin--dependent protein catabolic process
maturation of SSU-rRNA
sequence-specific mRNA binding
rRNA binding
structural constituent of ribosome
ligase activity
oxidoreductase activity
RNA binding
nucleotide binding
cytosolic small ribosomal subunit
cytosolic large ribosomal subunit
proteasome complex
ribosome
cytoplasmic stress granule
ribonucleoprotein complex
cytosol
Fold Enrichment
E
Comparison with literature
Aggregates in aged yeast Peters et al., 2012
SDS-resitant Chen et al., 2021
Palleted in stationary Narayanaswamy et al., 2009
Insoluble after 46°C HS Cherkasov et al, 2015
Extended SGs proteome Jain et al., 2016
Foci-forming at low pH Munder et al., 2016
P-body proteome Jain et al., 2016
SDS-resistant prion domains Alberti et al., 2009
No. of overlaping hits
-log(10) p.value.adj
F
%Cells with foci
Exponential
Stationary
Recovery
Asn1
Asn2
Bdh1
Gad1
Dcs1
G
%Cells with foci
0mM Potassium sorbate
8mM Potassium sorbate
H
%Cells with foci
0 min Adjusted to pH 7.5
30 min Adjusted to pH 7.5
0 min Non-adjusted pH
30 min Non-adjusted pH

**Supplementary Figure S3: Characterization of yeast proteins with predicted pH-sensing amyloid cores forming SDS-resistant and pH-dependent assemblies during stationary phase**

A. and B. Gene Ontology (GO) term enrichment among 1’523 human (A) and 576 yeast (B) proteins predicted to contain pH-sensing amyloid core motifs (Fig. 1F, 3A). Overrepresented biological process (BP, blue), molecular function (MF, orange), and cellular component (CC, grey) categories are shown.

C. Fractionation of 2 representative SDD–AGE gels separating proteins in cell extracts prepared from exponentially growing (Exp) or 3-day stationary phase (Stat) cells. Gel slices (F1 – F5) were identified using positions defined by the GeneRuler (GR) molecular weight marker visualized with Gel Red.

D. GO term enrichment of 551 proteins shifting in stationary phase to high-molecular weight assemblies in the SDD-AGE analysis (Fig. 3D). Overrepresented biological process (BP, blue), molecular function (MF, orange), and cellular component (CC, grey) categories are shown.

E. Yeast proteins shifting in stationary phase to high-molecular weight fractions in the SDD-AGE analysis (Fig. 3D) were compared with available literature data. Shown are hypergeometric test results with Holm p value correction, non-significant values are indicated in grey.

F. Quantification of microscopy data from Fig. 3E of cells expressing mNG-tagged Asn1, Asn2, Bdh1, Gad1, and Dcs1 during exponential growth, 2-day stationary phase, and after 2 h recovery from stationary phase. For each candidate, the percentage (%) of cells that form foci was determined; n=3, mean ± SD. A two-way ANOVA test, followed by Tukey’s multiple-comparisons, controls statistical significance; ** $p < 0.01$, **** $p < 0.0001$; ns, not significant.

G. Quantification of microscopy data from Fig. 3E of cells expressing mNG-tagged Asn1, Asn2, Bdh1, Gad1, and Dcs1 during exponential growth after 30 min treatment in the absence (0 mM) or presence (8 mM) of potassium sorbate to reduce intracellular pH. For each candidate, the percentage (%) of cells forming foci was determined; n=3. A two-way ANOVA test, followed by Tukey’s multiple-comparisons, controls statistical significance; **** $p < 0.0001$; only significant comparisons are shown.

H. Quantification of microscopy data from Fig. 3F of cells expressing mNG-tagged Asn1, Asn2, Bdh1, Gad1 and Dcs1 at 3-day stationary phase. The stationary-phase medium was then adjusted (pH 7.5) to increase intracellular pH or non-adjusted for control. For each candidate, the percentage (%) of cells with foci at time 0 and 30 min after media exchange was determined; n=3. A two-way ANOVA test, followed by Tukey's multiple-comparisons, controls statistical significance. * $p < 0.05$, ** $p < 0.01$, *** $p < 0.001$, **** $p < 0.0001$; ns, not significant.

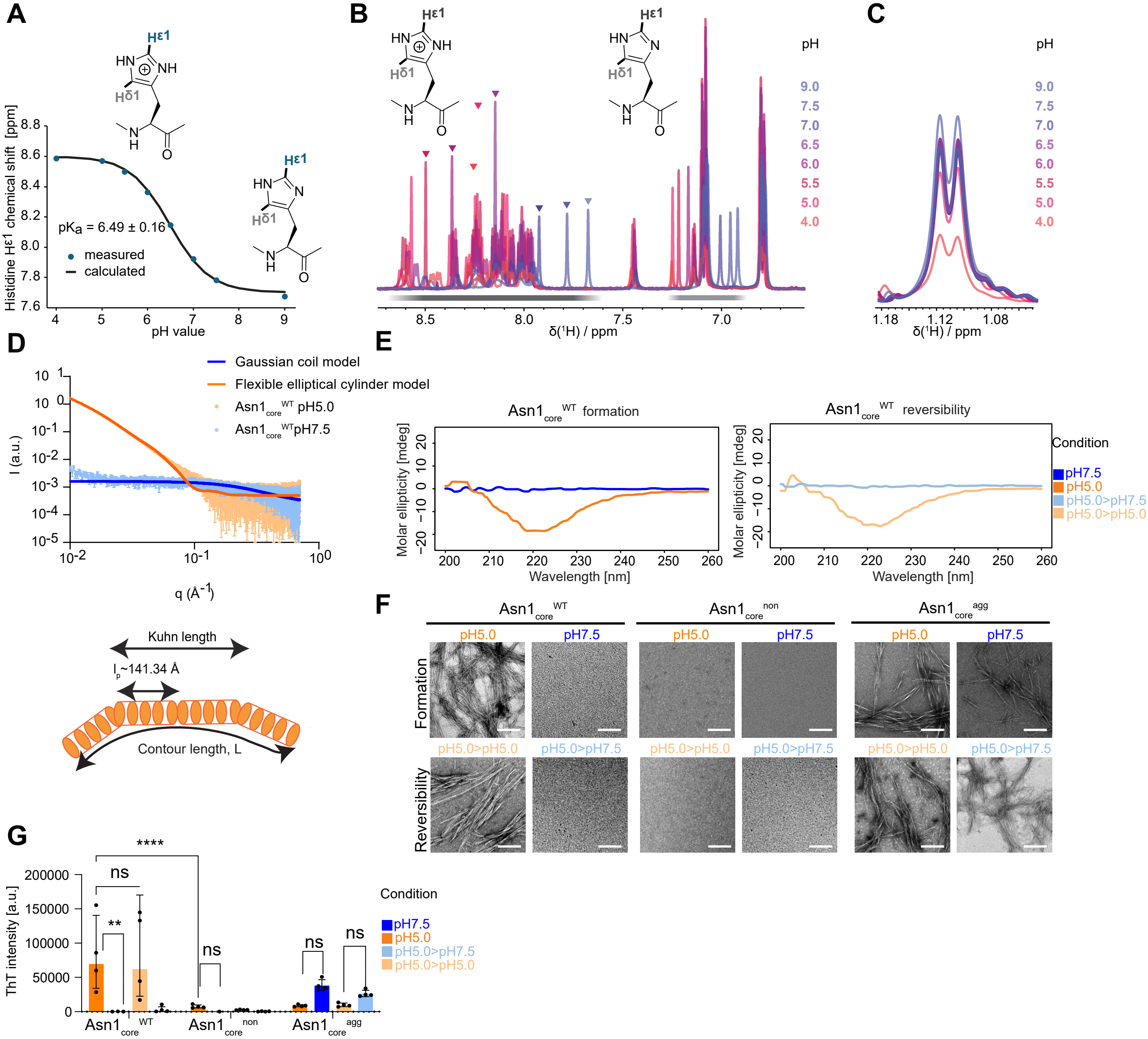
A
Hε1
HN
NH
Hδ1
N
H
O
Histidine Hε1 chemical shift [ppm]
pKa = 6.49 ± 0.16
measured
calculated
pH value
B
pH
9.0
7.5
7.0
6.5
6.0
5.5
5.0
4.0
δ(¹H) / ppm
C
δ(¹H) / ppm
D
Gaussian coil model
Flexible elliptical cylinder model
Asn1core WT pH5.0
Asn1core WT pH7.5
I (a.u.)
q (Å-1)
Kuhn length
lp~141.34 Å
Contour length, L
E
Asn1core WT formation
Asn1core WT reversibility
Molar ellipticity [mdeg]
Wavelength [nm]
Condition
pH7.5
pH5.0
pH5.0>pH7.5
pH5.0>pH5.0
F
Asn1core WT
Asn1core non
Asn1core agg
pH5.0
pH7.5
pH5.0>pH5.0
pH5.0>pH7.5
Formation
Reversibility
G
ThT intensity [a.u.]
****
ns
**
Condition
pH7.5
pH5.0
pH5.0>pH7.5
pH5.0>pH5.0
Asn1core WT
Asn1core non
Asn1core agg

**Supplementary Figure S4: pH-dependent Asn1 fibrillation is mediated by protonation of the central histidine in the predicted amyloid core**

A. NMR measurements revealed a pKa value of 6.49 ± 0.16 for the central histidine in the $Asn1_{core}^{WT}$ peptide, measured in a pH gradient from 4 to 9.

B. Overlay of 1D $^{1}H$ NMR spectra recorded across a pH gradient from 4 to 9, as indicated by the color code (right). The $^{1}H$ resonance corresponding to $H^{\varepsilon 1}$ of the central histidine within the $Asn1_{core}^{WT}$ peptide is indicated with triangles. The progressive chemical shift changes reflect protonation-dependent alterations of the histidine, which was used for the plot in panel A.

C. Excerpt of 1D $^{1}H$ NMR spectra of representative methyl resonances of the $Asn1_{core}^{WT}$ peptide recorded across a pH gradient from 4 to 9, as indicated by the color code (right). Disappearance of the signals reveals pH dependent aggregation of the peptide.

D. The structure of the $Asn1_{core}^{WT}$ peptide was analyzed by small-angle X-ray scattering (SAXS) (top) after 24 h incubation at 30°C at the indicated pH. SAXS data were visualized as log-log plot of the scattering intensity (I) versus the scattering vector (q). Schematic representation (bottom) of the modelled data at pH 5.0 shows a flexible elliptical cylinder. Data are representative of n=4 (dotted lines) and fitted to the corresponding models (solid lines).

E. Representative circular dichroism (CD) spectra of $Asn1_{core}^{WT}$ recorded after 24 h incubation at 30°C at pH 5.0 or pH 7.5 (formation, left). To probe reversibility, $Asn1_{core}^{WT}$ fibrils formed at pH 5.0 were analyzed after the pH was switched to 7.5 or kept at 5.0 for control for an additional 24 h at 30°C (right). Molar ellipticity (mdeg) as a function of wavelength is plotted; n=2.

F. Representative images of $Asn1_{core}^{WT}$, $Asn1_{core}^{non}$, $Asn1_{core}^{agg}$ peptides visualized by negative-staining TEM after 24 h incubation at pH 5.0 or pH 7.5 (formation). To probe reversibility, fibrils formed at pH 5.0 were imaged after the pH was switched to 7.5 or kept at 5.0 for control for an additional 24 h at 30°C; n=3, scale bar: 200 nm.

G. ThT binding of $Asn1_{core}^{WT}$, $Asn1_{core}^{non}$ and $Asn1_{core}^{agg}$ peptides measured after 24 h incubation at 30°C pH 5.0 or pH 7.5 (formation). To probe reversibility, fibrils formed at pH 5.0 were analyzed after the pH was switched to 7.5 or kept at 5.0 for control for an additional 24 h at 30°C; n= 4, mean ± SD; a.u.: arbitrary units. A two-way ANOVA test, followed by Tukey's multiple-comparisons, controls statistical significance; ** $p < 0.01$, **** $p < 0.0001$; ns, not significant; only relevant comparisons are shown.

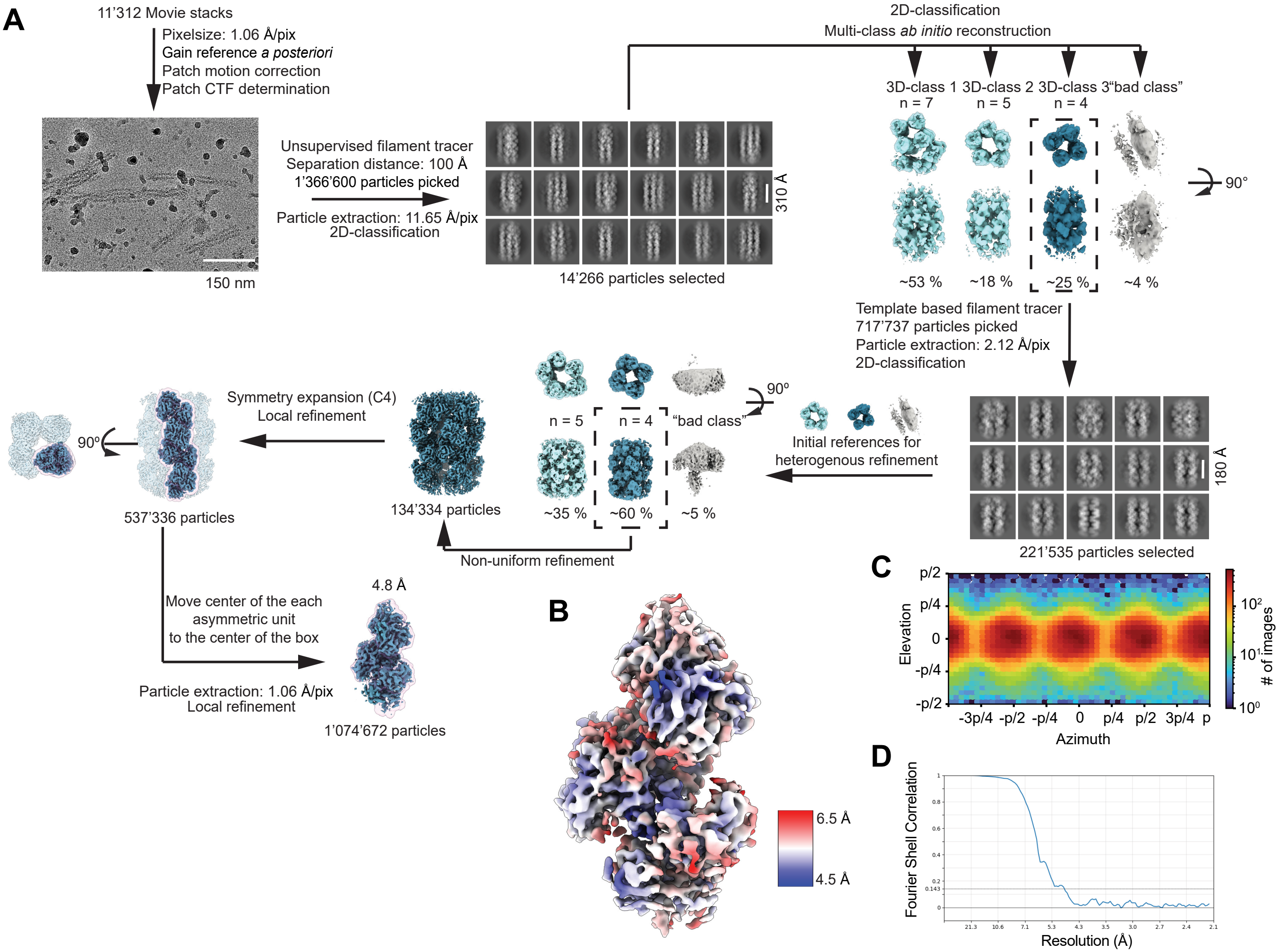
A
11'312 Movie stacks
Pixelsize: 1.06 Å/pix
Gain reference a posteriori
Patch motion correction
Patch CTF determination
150 nm
Unsupervised filament tracer
Separation distance: 100 Å
1'366'600 particles picked
Particle extraction: 11.65 Å/pix
2D-classification
310 Å
14'266 particles selected
2D-classification
Multi-class ab initio reconstruction
3D-class 1 3D-class 2 3D-class 3"bad class"
n = 7
n = 5
n = 4
90°
~53 %
~18 %
~25 %
~4 %
Template based filament tracer
717'737 particles picked
Particle extraction: 2.12 Å/pix
2D-classification
180 Å
221'535 particles selected
Initial references for
heterogenous refinement
90°
n = 5
n = 4
"bad class"
~35 %
~60 %
~5 %
Non-uniform refinement
134'334 particles
Symmetry expansion (C4)
Local refinement
90°
537'336 particles
Move center of the each
asymmetric unit
to the center of the box
Particle extraction: 1.06 Å/pix
Local refinement
4.8 Å
1'074'672 particles
B
6.5 Å
4.5 Å
C
p/2
p/4
0
-p/4
-p/2
Elevation
-3p/4 -p/2 -p/4 0 p/4 p/2 3p/4 p
Azimuth
10²
10¹
10⁰
### of images
D
Fourier Shell Correlation
Resolution (Å)

**Supplementary Figure S5. Single-particle analysis of Asn1$_{FL}$$^{WT}$ shows fibril formation at low pH**

A. Cryo-EM single-particle analysis workflow in cryoSPARC (Punjani *et al.*, 2017).
B. Final cryo-EM density map of Asn1$_{FL}$$^{WT}$ fibril colored by local resolution estimation.
C. Euler angle distribution of particles contributing to the final reconstruction.
D. Fourier shell correlation (FSC) curve indicating an overall resolution of 4.8 Å.

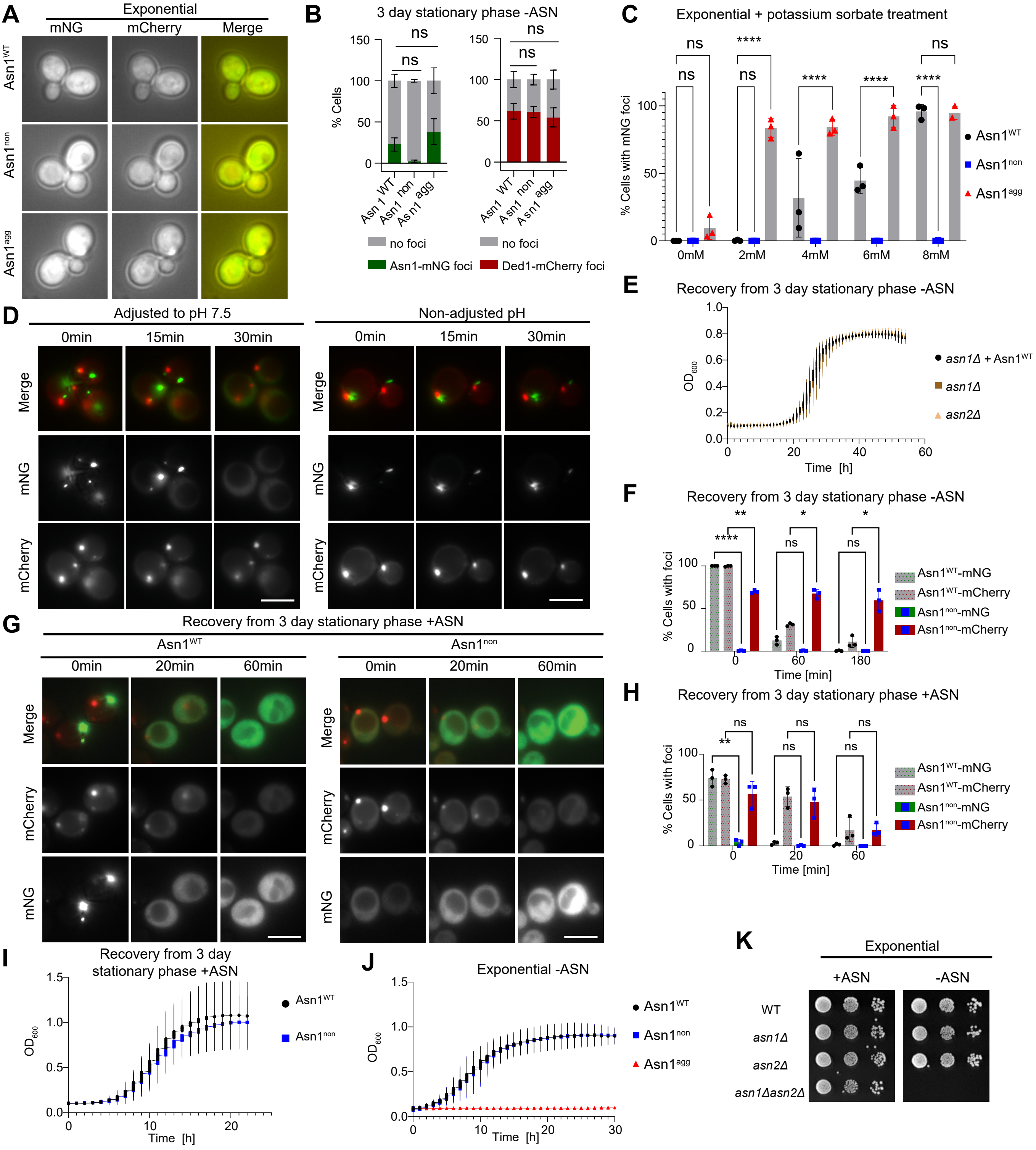
A
Exponential
mNG
mCherry
Merge
Asn1^WT
Asn1^non
Asn1^agg
B
3 day stationary phase -ASN
ns
ns
% Cells
150
100
50
0
Asn 1 WT
Asn1 non
Asn1 agg
no foci
Asn1-mNG foci
Ded1-mCherry foci
C
Exponential + potassium sorbate treatment
****
% Cells with mNG foci
0mM
2mM
4mM
6mM
8mM
Asn1^WT
Asn1^non
Asn1^agg
D
Adjusted to pH 7.5
Non-adjusted pH
0min
15min
30min
Merge
mNG
mCherry
E
Recovery from 3 day stationary phase -ASN
OD600
Time [h]
asn1Δ + Asn1^WT
asn1Δ
asn2Δ
F
Recovery from 3 day stationary phase -ASN
% Cells with foci
Time [min]
Asn1^WT-mNG
Asn1^WT-mCherry
Asn1^non-mNG
Asn1^non-mCherry
G
Recovery from 3 day stationary phase +ASN
Asn1^WT
Asn1^non
0min
20min
60min
H
Recovery from 3 day stationary phase +ASN
I
Recovery from 3 day stationary phase +ASN
J
Exponential -ASN
K
Exponential
+ASN
-ASN
WT
asn1Δ
asn2Δ
asn1Δasn2Δ

**Supplementary Figure S6. *In vivo* characterization of pH-dependent Asn1 assembly**

A. *asn1*Δ cells harboring the SG marker Ded1-mCherry and stably expressing mNG-tagged $Asn1^{WT}$, $Asn1^{non}$ or $Asn1^{agg}$ from the endogenous promoter were imaged during exponential growth in complete media; representative images, n=3, scale bar: 5 µm.

B. Quantification of foci-containing *asn1*Δ cells harboring Ded1-mCherry and stably expressing mNG-tagged $Asn1^{WT}$, $Asn1^{non}$ or $Asn1^{agg}$ from the endogenous promoter imaged after 3 days stationary phase in minimal medium without asparagine (-ASN). A two-way ANOVA followed by Dunnett's multiple-comparisons test controls statistical significance vs. WT, n = 3, mean ± SD; ns: not significant; scale bar: 5 µm.

C. Quantification of foci-containing *asn1*Δ cells harboring the SG marker Ded1-mCherry and stably expressing mNG-tagged $Asn1^{WT}$, $Asn1^{non}$ or $Asn1^{agg}$ from the endogenous promoter imaged during exponential growth in complete medium and treated with 0, 2, 4, 6 or 8 mM potassium sorbate for 30 min to decrease intracellular pH; n=3, mean ± SD. A two-way ANOVA followed by Dunnett's multiple-comparisons test controls statistical significance; **** $p < 0.0001$ vs. WT; ns: not significant; scale bar: 5 µm.

D. *asn1*Δ cells stably expressing mNG-tagged $Asn1^{WT}$ and Ded1-mCherry were imaged during 3-day stationary phase in complete media, following a change to the stationary phase media adjusted to high pH (pH 7.5) or kept non-adjusted for control; representative images, n=3, scale bar: 5 µm. Note that increasing intracellular pH in stationary phase cells is sufficient to dissolve Asn1 foci but not SGs.

E. Recovery of *asn1*Δ cells stably expressing from the endogenous promoter mNG-tagged $Asn1^{WT}$, or *asn1*Δ or *asn2*Δ cells from 3-day stationary phase after diluting in fresh minimal media lacking asparagine (-ASN) was monitored by optical density ($OD_{600}$) measurements; n=3, mean ± SD.

F. Quantification of data in Fig. 5D. 3-day stationary phase *asn1*Δ cells harboring Ded1-mCherry and stably expressing mNG-tagged $Asn1^{WT}$ or $Asn1^{non}$ from the endogenous promoter (time 0), followed by their recovery at 60 and 180 min after addition of fresh minimal media without asparagine (-ASN). Cells with $Asn1^{WT}$ foci were selected and tracked over time (WT t(0) = 100%). The percentage (%) of cells with foci were counted at each time point; n=3, mean ± SD. A two-way ANOVA followed by Tukey's multiple-comparisons test controls statistical significance. * $p < 0.05$, ** $p < 0.01$, **** $p < 0.0001$; ns, not significant.

G. *asn1*Δ cells harboring Ded1-mCherry and stably expressing mNG-tagged Asn1$^{WT}$ or Asn1$^{non}$ from the endogenous promoter were imaged after 3 days in stationary phase (time 0) in minimal medium with asparagine (+ASN). Fresh minimal medium (+ASN) was then added and recovery monitored after 20 and 60 min; representative images, n=3, scale bar: 5 µm.

H. Quantification of data in panel G. The percentage (%) of cells with foci were counted at each time point; n=3, mean ± SD. A two-way ANOVA followed by Tukey's multiple-comparisons test controls statistical significance. ** $p < 0.01$, ns: not significant.

I. After 3-day stationary phase in minimal media with asparagine (+ASN) *asn1*Δ cells stably expressing Asn1$^{WT}$ or Asn1$^{non}$ from the endogenous promoter were diluted into fresh medium (+ASN), and the recovery was monitored by optical density ($OD_{600}$) measurements; n=3, mean ± SD.

J. Exponential growth of freshly diluted *asn1*Δ *asn2*Δ cells stably expressing mNG-tagged Asn1$^{WT}$, Asn1$^{non}$ or Asn1$^{agg}$ from the endogenous promoter was monitored by optical density ($OD_{600}$) measurements in minimal medium without asparagine (-ASN); n=3, mean ±SD.

K. Representative serial dilution spottings of exponentially growing wild-type, *asn1*Δ, *asn2*Δ and *asn1*Δ *asn2*Δ cells on minimal media with (+ASN) or without asparagine (-ASN); n=3. Plates were photographed after 3 days at 30$^{0}$C.

**Supplementary Table S1. Peptides tested in this study.** Table provided in a separate Excel file; contains all peptides except for the TAPAAS-tested ones.

**Supplementary Table S2. TAPAAS peptides.** Table provided in a separate Excel file.

**Supplementary Table S3. Summary of the TAPAAS screen results in respect to the initial bioinformatic predictions.** Number of peptides within given categories are presented. The prediction performance was calculated only based on the unambiguously classified peptides (n=309). Shaded in green are the correct predictions, resulting in accuracy of 34%. The true positive rate (TPR; proportion of true positives to all positives) was 75%, false positive rate (FPR; proportion of false positives to all negatives) was 64% (or specifically 42% for non-amyloids and 81% for constitutive amyloids. The precision was defined as true positives to all predicted positives (n=201, excluding the predictions on ambiguous and excluded peptides) was approximately 10%.

<table>
<tr><th colspan="2" rowspan="2">Prediction</th><th colspan="5">TAPAAS outcome</th></tr>
<tr><th colspan="3">unambiguously classified:</th><th rowspan="2">ambiguous</th><th rowspan="2">excluded</th></tr>
<tr><th>class</th><th>n</th><th>pH-sensing</th><th>non-amyloid</th><th>constitutive</th></tr>
<tr><td>pH-sensing</td><td>306</td><td>21</td><td>53</td><td>127</td><td>76</td><td>29</td></tr>
<tr><td>non-amyloid</td><td>176</td><td>6</td><td>70</td><td>16</td><td>48</td><td>36</td></tr>
<tr><td>constitutive</td><td>22</td><td>1</td><td>2</td><td>13</td><td>4</td><td>2</td></tr>
<tr><td rowspan="2">total</td><td rowspan="2">504</td><td colspan="3">309:</td><td rowspan="2">128</td><td rowspan="2">67</td></tr>
<tr><td>28</td><td>125</td><td>156</td></tr>
</table>

**Supplementary Table S4: Bioinformatic predictions.** Table provided in a separate Excel file.

**Supplementary Table S5: Summary of the performance of the improved predictions on the TAPAAS screen results.** Number of peptides within given categories are presented. The prediction performance was calculated only based on the 309 unambiguously classified peptides. Shaded in green are the correct predictions, resulting in accuracy of 65.7%. TPR was 78.6%, FPR was 15.7% (21.6% for non-amyloids; 10.9% for constitutive amyloids). The precision was 33.3%.

| Prediction | | TAPAAS outcome | | |
|---|---|---|---|---|
| **class** | **n** | **pH-sensing** | **non-amyloid** | **constitutive** |
| **pH-sensing** | 66 | 22 | 27 | 17 |
| **non-amyloid** | 80 | 2 | 60 | 18 |
| **constitutive** | 163 | 4 | 38 | 121 |
| **total** | 309 | 28 | 125 | 156 |

**Supplementary Table S6: Fitting and modeling the SAXS data into SasView.**

| **pH** | **Sample** | **Model** | **$R_g$ (Å)** | **$I_0$ ($cm^{-1}$)** | **$\chi^2/N_{pts}$** | |
|---|---|---|---|---|---|---|
| 7.5 | $Asn1_{core}^{WT}$ | Mono Gaussian coil model | 6.68±0.33 | 0.0020±0.00012 | 0.068 | |
| 7.5 | ARA1 | Mono Gaussian coil model | 7.76±0.41 | 0.0016±0.0001 | 0.015 | |
| **pH** | **Sample** | **Model** | **Kuhn Length (Å)** | **Radius (Å)** | **Axis Ratio** | **χ2/Npts** |
| 5.0 | $Asn1_{core}^{WT}$ | Flexible elliptical cylinder | 141.34±1.56 | 34.21±0.13 | 3.61±0.01 | 0.8 |
| 5.0 | ARA1 | Cylinder elliptical | Used a fixed cylinder length of 10000 Å | 29.37±0.04 | 3.58±0.01 | 0.98 |

**Supplementary Table S7: GO terms enrichment analysis.** Table provided in a separate Excel file.

**Supplementary Table S8: Fluorescence microscopy screen on yeast candidate proteins.** Table provided in a separate Excel file.

**Supplementary Table S9: SDD-AGE-MS results.** Table provided in a separate Excel file.

**Supplementary Table S10: Plasmids used in this study.**

| Catalog ID | Name | Relevant features | Origin |
|---|---|---|---|
| pJK476 | pSMT3_scASN1-6xHis | Purification plasmid for $Asn1_{FL}^{WT}$ | Gift from Kollman laboratory |
| AK344 | pSMT3_scASN1-H334A-6xHis | Purification plasmid for $Asn1_{FL}^{non}$ | This study |
| AK346 | pSMT3_scASN1-D329N, D330N, D339N, E336Q-6xHis | Purification plasmid for $Asn1_{FL}^{agg}$ | This study |
| AK7 | pFA6a-mNeonGreen-KanMX6 | Deletion Kan | Addgene |
| AK8 | pFA6a-mNeonGreen-HIS3MX6 | mNG tagging | Addgene |

**Supplementary Table S11: Data collection, 3D reconstruction and model statistics.**

| | | $\mathbf{Asn1_{FL}^{WT}}$ |
|---|---|---|
| **Data collection** | | |
| Micrographs | | 11’312 |
| Microscope Voltage (kV) | | 300 |
| Detector | K3 camera | |
| Pixel size (Å/pixel) | | 1.06 |
| Target defocus range (μm) | | -1.2 to -2.8 |
| Total dose (e-/Å²) | | 48 |
| Software | EPU (v. 3.10.0) | |
| **Reconstruction** | | |
| Software | cryoSPARC (v. 5.0.2) | |
| Final particles (No.) | | 1'074’672 |
| Resolution, 0.143 threshold (Å) | | 4.8 |
| Symmetry imposed | C4 | |
| Map sharpening B-factor (Å²) | | -345 |
| **Model building** | | |
| Initial model used | AlphaFold3 | |
| Software | Coot (v. 9.8.5) | |
| Residues | | 1’144 |
| Non-hydrogen atoms | | 9’084 |
| **Refinement** | | |
| Software | Phenix (v. 1.21.2) | |
| R.M.S deviations | | |
| Bond length (Å) | | 0.002 |
| Bond angles (°) | | 0.512 |
| Validation | | |
| MolProbity score | | 1.45 |
| Clashscore | | 6.73 |
| Rotamer outliers (%) | | 0.00 |
| Ramachandran plot | | |
| Favored (%) | | 96.67 |
| Allowed (%) | | 2.37 |
| Outliers (%) | | 0.00 |

**Supplementary Table S12: Yeast strains used in this study.** *GFP collection - Thermo Fisher Scientific, (Huh *et al.*, 2003)

| Strain ID | Name | Genotype | Origin |
|---|---|---|---|
| yCW1110 | Asn1D:KAN Asn2D:NAT | BY4741, asn1Δ::KANR; asn2Δ::NATR | This study |
| yCW1111 | Asn1-mNG:URA Asn1D:KAN Asn2D:NAT | BY4741, asn1Δ::KANR; Asn1-mNG::URA; asn2Δ::NATR | This study |
| yCW1112 | Asn1non-mNG:URA Asn1D:KAN Asn2D:NAT | BY4741, asn1Δ::KANR; Asn1-H334A-mNG::URA; asn2Δ::NATR | This study |
| yCW1115 | Asn1agg-mNG:URA Asn1D:KAN Asn2D:NAT | BY4741, asn1Δ::KANR; Asn1-D329N, D330N, D339N, E336Q-mNG::URA; asn2Δ::NATR | This study |
| yCW1116 | Asn1-mNG:URA Ded1-cherry:KAN asn1D:KAN | BY4741, asn1Δ::KANR; Asn1-mNG::URA; DED1::DED1-mCherry::KAN$^{R}$ | This study |
| yCW1117 | Asn1-mNG:URA Ded1-cherry:KAN asn1D:KAN | BY4742, asn1Δ::KANR; Asn1-mNG::URA; DED1::DED1-mCherry::KAN$^{R}$ | This study |
| yCW1120 | Asn1agg-mNG:URA Ded1-cherry:KAN asn1D:KAN | BY4742, asn1Δ::KANR; Asn1-D329N, D330N, D339N, E336Q-mNG::URA; DED1::DED1-mCherry::KAN$^{R}$ | This study |
| yAK121 | asn1D:KAN | BY4741, asn1Δ::KANR | This study |
| yAK238 | asn2D:KAN | BY4741, asn2Δ::KANR | This study |
| yCW1135 | Asn1non-mNG:URA Asn1D:KAN Ded1:KAN | BY4741, asn1Δ::KANR; Asn1-H334A-mNG::URA; DED1::DED1-mCherry::KAN$^{R}$ | This study |
| yAK129 | Asn1-mNg:HIS | BY4741, Asn1-mNg:HIS | This study |
| yAK130 | Asn2-mNg:HIS | BY4741, Asn2-mNg:HIS | This study |
| yAK387 | Dcs1-mNg:HIS | BY4741, Dcs1-mNg:HIS | This study |
| yAK392 | Bdh1mNg:HIS | BY4741, Bdh1-mNg:HIS | This study |
| yAK393 | Gad1-mNg:HIS | BY4741, Gad1-mNg:HIS | This study |
| COL87430 | Utp10-GFP:HIS | BY4741, Utp10-GFP:HIS | GFP collection* |
| COL87470 | Iqg1-GFP:HIS | BY4741, Iqg1-GFP:HIS | GFP collection* |
| COL87499 | Bem2-GFP:HIS | BY4741, Bem2-GFP:HIS | GFP collection* |
| COL87450 | Tom1-GFP:HIS | BY4741, Tom1-GFP:HIS | GFP collection* |
| COL87456 | Uls1-GFP:HIS | BY4741, Uls1-GFP:HIS | GFP collection* |
| COL87460 | Trm3-GFP:HIS | BY4741, Trm3-GFP:HIS | GFP collection* |
| COL87439 | Acc1-GFP:HIS | BY4741, Acc1-GFP:HIS | GFP collection* |
| COL87465 | Ubr2-GFP:HIS | BY4741, Ubr2-GFP:HIS | GFP collection* |
| COL87513 | Rim15-GFP:HIS | BY4741, Rim15-GFP:HIS | GFP collection* |
| COL87482 | Scc2-GFP:HIS | BY4741, Scc2-GFP:HIS | GFP collection* |
| COL87480 | Atg2-GFP:HIS | BY4741, Atg2-GFP:HIS | GFP collection* |
| COL87527 | Tda9-GFP:HIS | BY4741, Tda9-GFP:HIS | GFP collection* |
| COL87542 | Atg26-GFP:HIS | BY4741, Atg26-GFP:HIS | GFP collection* |
| COL87609 | Pds5-GFP:HIS | BY4741, Pds5-GFP:HIS | GFP collection* |
| COL87540 | Smc3-GFP:HIS | BY4741, Smc3-GFP:HIS | GFP collection* |
| COL87612 | Smc1-GFP:HIS | BY4741, Smc1-GFP:HIS | GFP collection* |
| COL87601 | Axl1-GFP:HIS | BY4741, Axl1-GFP:HIS | GFP collection* |
| COL87543 | Inp52-GFP:HIS | BY4741, Inp52-GFP:HIS | GFP collection* |

| | | | |
|---|---|---|---|
| COL87606 | Rse1-GFP:HIS | BY4741, Rse1-GFP:HIS | GFP collection* |
| COL87559 | Cft1-GFP:HIS | BY4741, Cft1-GFP:HIS | GFP collection* |
| COL87616 | Irr1-GFP:HIS | BY4741, Irr1-GFP:HIS | GFP collection* |
| COL87657 | Psk2-GFP:HIS | BY4741, Psk2-GFP:HIS | GFP collection* |
| COL87671 | Hos4-GFP:HIS | BY4741, Hos4-GFP:HIS | GFP collection* |
| COL87638 | Pik1-GFP:HIS | BY4741, Pik1-GFP:HIS | GFP collection* |
| COL87660 | Hrq1-GFP:HIS | BY4741, Hrq1-GFP:HIS | GFP collection* |
| COL87633 | Bud2-GFP:HIS | BY4741, Bud2-GFP:HIS | GFP collection* |
| COL87705 | Pol3-GFP:HIS | BY4741, Pol3-GFP:HIS | GFP collection* |
| COL87689 | Met18-GFP:HIS | BY4741, Met18-GFP:HIS | GFP collection* |
| COL87659 | Crm1-GFP:HIS | BY4741, Crm1-GFP:HIS | GFP collection* |
| COL87632 | Pan2-GFP:HIS | BY4741, Pan2-GFP:HIS | GFP collection* |
| COL87795 | Utp21-GFP:HIS | BY4741, Utp21-GFP:HIS | GFP collection* |
| COL87804 | Vid30-GFP:HIS | BY4741, Vid30-GFP:HIS | GFP collection* |
| COL87715 | Lrg1-GFP:HIS | BY4741, Lrg1-GFP:HIS | GFP collection* |
| COL87806 | Sxm1-GFP:HIS | BY4741, Sxm1-GFP:HIS | GFP collection* |
| COL87731 | Hsh155-GFP:HIS | BY4741, Hsh155-GFP:HIS | GFP collection* |
| COL87739 | Gpi13-GFP:HIS | BY4741, Gpi13-GFP:HIS | GFP collection* |
| COL87832 | Psy2-GFP:HIS | BY4741, Psy2-GFP:HIS | GFP collection* |
| COL87888 | Dug2-GFP:HIS | BY4741, Dug2-GFP:HIS | GFP collection* |
| COL87869 | Spc98-GFP:HIS | BY4741, Spc98-GFP:HIS | GFP collection* |
| COL87859 | Pms1-GFP:HIS | BY4741, Pms1-GFP:HIS | GFP collection* |
| COL87835 | Sap155-GFP:HIS | BY4741, Sap155-GFP:HIS | GFP collection* |
| COL87853 | Hir2-GFP:HIS | BY4741, Hir2-GFP:HIS | GFP collection* |
| COL87926 | Kip3-GFP:HIS | BY4741, Kip3-GFP:HIS | GFP collection* |
| COL87930 | Rtt101-GFP:HIS | BY4741, Rtt101-GFP:HIS | GFP collection* |
| COL88048 | Sky1-GFP:HIS | BY4741, Sky1-GFP:HIS | GFP collection* |
| COL88059 | Gyp7-GFP:HIS | BY4741, Gyp7-GFP:HIS | GFP collection* |
| COL88069 | Gcn20-GFP:HIS | BY4741, Gcn20-GFP:HIS | GFP collection* |
| COL88012 | Cul3-GFP:HIS | BY4741, Cul3-GFP:HIS | GFP collection* |
| COL88066 | Avl9-GFP:HIS | BY4741, Avl9-GFP:HIS | GFP collection* |
| COL88053 | Prp43-GFP:HIS | BY4741, Prp43-GFP:HIS | GFP collection* |
| COL88023 | Nto1-GFP:HIS | BY4741, Nto1-GFP:HIS | GFP collection* |
| COL88179 | Hos3-GFP:HIS | BY4741, Hos3-GFP:HIS | GFP collection* |
| COL88116 | Spp382-GFP:HIS | BY4741, Spp382-GFP:HIS | GFP collection* |
| COL88121 | Sse1-GFP:HIS | BY4741, Sse1-GFP:HIS | GFP collection* |
| COL88155 | Trs85-GFP:HIS | BY4741, Trs85-GFP:HIS | GFP collection* |
| COL88224 | Chs5-GFP:HIS | BY4741, Chs5-GFP:HIS | GFP collection* |
| COL88273 | Ufo1-GFP:HIS | BY4741, Ufo1-GFP:HIS | GFP collection* |
| COL88361 | Vps27-GFP:HIS | BY4741, Vps27-GFP:HIS | GFP collection* |
| COL88296 | Uba2-GFP:HIS | BY4741, Uba2-GFP:HIS | GFP collection* |
| COL88468 | Gad1-GFP:HIS | BY4741, Gad1-GFP:HIS | GFP collection* |
| COL88497 | Yeh1-GFP:HIS | BY4741, Yeh1-GFP:HIS | GFP collection* |

| | | | |
|---|---|---|---|
| COL88486 | Asn1-GFP:HIS | BY4741, Asn1-GFP:HIS | GFP collection* |
| COL88545 | Asn2-GFP:HIS | BY4741, Asn2-GFP:HIS | GFP collection* |
| COL88742 | Pyk2-GFP:HIS | BY4741, Pyk2-GFP:HIS | GFP collection* |
| COL88771 | Ubp6-GFP:HIS | BY4741, Ubp6-GFP:HIS | GFP collection* |
| COL88830 | Ald6-GFP:HIS | BY4741, Ald6-GFP:HIS | GFP collection* |
| COL88893 | Rnt1-GFP:HIS | BY4741, Rnt1-GFP:HIS | GFP collection* |
| COL88917 | YMR027W-GFP:HIS | BY4741, YMR027W-GFP:HIS | GFP collection* |
| COL89027 | Osh6-GFP:HIS | BY4741, Osh6-GFP:HIS | GFP collection* |
| COL89046 | Cdc40-GFP:HIS | BY4741, Cdc40-GFP:HIS | GFP collection* |
| COL89005 | Rpn5-GFP:HIS | BY4741, Rpn5-GFP:HIS | GFP collection* |
| COL89025 | Fbp26-GFP:HIS | BY4741, Fbp26-GFP:HIS | GFP collection* |
| COL88991 | Sah1-GFP:HIS | BY4741, Sah1-GFP:HIS | GFP collection* |
| COL89059 | Hog1-GFP:HIS | BY4741, Hog1-GFP:HIS | GFP collection* |
| COL89189 | Dcs2-GFP:HIS | BY4741, Dcs2-GFP:HIS | GFP collection* |
| COL89318 | Bdh1-GFP:HIS | BY4741, Bdh1-GFP:HIS | GFP collection* |
| COL89350 | Lys12-GFP:HIS | BY4741, Lys12-GFP:HIS | GFP collection* |
| COL89448 | Dcs1-GFP:HIS | BY4741, Dcs1-GFP:HIS | GFP collection* |
| COL89525 | Mob1-GFP:HIS | BY4741, Mob1-GFP:HIS | GFP collection* |
| COL89613 | Sds22-GFP:HIS | BY4741, Sds22-GFP:HIS | GFP collection* |
| COL89539 | Gtr2-GFP:HIS | BY4741, Gtr2-GFP:HIS | GFP collection* |
| COL89718 | Ect1-GFP:HIS | BY4741, Ect1-GFP:HIS | GFP collection* |
| COL89795 | Rme1-GFP:HIS | BY4741, Rme1-GFP:HIS | GFP collection* |
| COL89951 | Mam33-GFP:HIS | BY4741, Mam33-GFP:HIS | GFP collection* |
| COL90047 | Pre8-GFP:HIS | BY4741, Pre8-GFP:HIS | GFP collection* |
| COL90163 | YPL199C-GFP:HIS | BY4741, YPL199C-GFP:HIS | GFP collection* |
| COL90857 | Nup192-GFP:HIS | BY4741, Nup192-GFP:HIS | GFP collection* |
| COL90880 | Cyr1-GFP:HIS | BY4741, Cyr1-GFP:HIS | GFP collection* |
| COL90903 | Sec26-GFP:HIS | BY4741, Sec26-GFP:HIS | GFP collection* |
| COL90904 | Cwh43-GFP:HIS | BY4741, Cwh43-GFP:HIS | GFP collection* |
| COL90962 | Vps41-GFP:HIS | BY4741, Vps41-GFP:HIS | GFP collection* |
| COL90925 | Vam6-GFP:HIS | BY4741, Vam6-GFP:HIS | GFP collection* |
| COL90887 | Neo1-GFP:HIS | BY4741, Neo1-GFP:HIS | GFP collection* |
| COL91143 | Ret2-GFP:HIS | BY4741, Ret2-GFP:HIS | GFP collection* |
| COL91101 | Rli1-GFP:HIS | BY4741, Rli1-GFP:HIS | GFP collection* |
| COL91203 | Cot1-GFP:HIS | BY4741, Cot1-GFP:HIS | GFP collection* |
| COL91477 | Frq1-GFP:HIS | BY4741, Frq1-GFP:HIS | GFP collection* |
